# Pre-existing chromatin accessibility primes δ-cells for injury-induced endocrine plasticity

**DOI:** 10.64898/2026.05.24.727280

**Authors:** Jo Gräßlin, Prateek Chawla, Padmapriya Subramanian, Anuradha Rajendran, Emilie Ida Christine Walda, Alexander Froschauer, Nikolay Ninov, Jan Philipp Junker

## Abstract

Adult zebrafish rapidly recover glucose homeostasis after β-cell loss, but the cellular basis and regulatory mechanisms that enable this response remain unclear. Here we combine single-cell transcriptomics, single-cell chromatin accessibility profiling, paired multiome analysis and functional perturbation to define early pancreatic recovery after β-cell ablation. We show that, during the first month after injury, insulin production is restored predominantly by *sst1.1*^+^ δ1-cells rather than by rapid reconstitution of canonical β-cells. Following ablation, δ1-cells adopt a bihormonal hybrid state and induce metabolic, secretory and β-cell-associated gene programs. Systematic comparison of chromatin accessibility across endocrine cell types reveals that these δ1-cells are uniquely close to β-cells and exhibit open chromatin at β-cell enhancers in the steady state. Moreover, hybrid-cell formation occurs without major chromatin remodeling, with β-cell associated loci being already accessible in δ1-cells before β-cell injury. A comparable permissive state is present in medaka but not in human δ-cells, suggesting that restricted insulin accessibility may represent a barrier to endocrine plasticity in the human pancreas. In zebrafish, δ1-cells also show evidence of metabolic remodeling after β-cell loss, including rapid accumulation of neutral lipids. Gene regulatory network analysis and perturbation identify *meis1a/b* as required regulators of δ1 hybrid-cell formation after β-cell loss in zebrafish, whereas *meis1a/b* are not upregulated following β-cell ablation in medaka. Together, our results define pre-existing chromatin accessibility, metabolic remodeling and instructive transcriptional regulation as key features of early functional recovery after β-cell loss in the adult zebrafish pancreas.

## Introduction

Loss or dysfunction of insulin-producing pancreatic β-cells is a central driver of diabetes mellitus. Although modern insulin replacement and glucose-lowering therapies have transformed disease management, they do not restore the endogenous regulatory logic of the islet and therefore cannot fully prevent long-term complications. A major goal in regenerative and translational biology is to replace β-cell function durably – either by stimulating β-cell regeneration *in situ* or by inducing alternative endocrine cell types to acquire β-like properties (Karampelias *et al*, 2025; Aguayo-Mazzucato & Bonner-Weir, 2018). Progress toward this goal has been limited by an incomplete understanding of which adult cell states can execute β-cell replacement programs and what regulatory constraints prevent such fate changes in species with low regenerative capacity.

Among vertebrate models, zebrafish stand out for their ability to rapidly recover glucose homeostasis after profound β-cell loss in the adult stage (Moss *et al*, 2009). This regenerative competence has made zebrafish a powerful system to discover cellular behaviors that are difficult to elicit or observe in mammals. Insulin-positive cells reappear after injury and have variously been interpreted as newly formed β-cells arising through proliferation, neogenesis from non-endocrine compartments, or transdifferentiation of related endocrine populations (Moss *et al*, 2009; Delaspre *et al*, 2015; Carril Pardo *et al*, 2022; Singh *et al*, 2022; Mi *et al*, 2023). At the same time, increasing evidence across vertebrates suggests that endocrine identity is not strictly discrete: polyhormonal and “hybrid” endocrine states exist during development and can emerge in response to stress or injury (Bocian-Sobkowska *et al*, 1999; Chera *et al*, 2014; Singh *et al*, 2022; Carril Pardo *et al*, 2022). In this context, reliance on insulin expression alone as a marker can blur the distinction between bona fide β-cells and other endocrine cell types that transiently (or stably) activate insulin as part of an adaptive response.

A major challenge, therefore, is to define – unbiasedly and with sufficient resolution – which cells actually provide insulin during functional recovery, and whether different proposed sources represent parallel, sequential, or context-dependent routes. Equally important is to identify the regulatory mechanisms that enable these transitions. Cell identity changes typically require not only instructive signals and transcription factor activity, but also a permissive chromatin landscape that allows the activation of the respective lineage programs that are normally silenced. Such epigenetic constraints can act as “roadblocks” to plasticity, potentially explaining why robust β-cell replacement is achievable in some species and injury paradigms but limited in adult mammals (Puri *et al*, 2015; Arda *et al*, 2018; Sun *et al*, 2021). An integrated view of transcriptional state, regulatory potential, and cell-type heterogeneity is thus needed to understand the logic of endocrine plasticity in vivo and to pinpoint leverage points for enhancing regeneration.

In zebrafish, the response to β-cell loss can be probed with temporal control using targeted ablation strategies such as the nitroreductase (NTR) system, in which NTR expression driven by the insulin promoter enables selective elimination of β-cells upon metronidazole (MTZ) treatment (Figure 1A) (Curado *et al*, 2008; Tucker *et al*, 2023). This approach creates a strong physiological demand for insulin and provides a defined time window in which to follow endocrine adaptation. Here, we use β-cell ablation in zebrafish, combined with comparative analysis in medaka and human pancreas, to define how injury-induced endocrine plasticity is enabled or constrained. We show that insulin restoration is executed by a primed δ1-cell state, identify a permissive chromatin landscape that precedes injury, uncover metabolic remodeling as a functional component of the response, and define meis1a/b as instructive regulators of insulin induction. These findings distinguish permissive, deployable, and blocked states of endocrine plasticity and provide a general framework for understanding fate switching in differentiated adult tissues.

**Figure 1:**
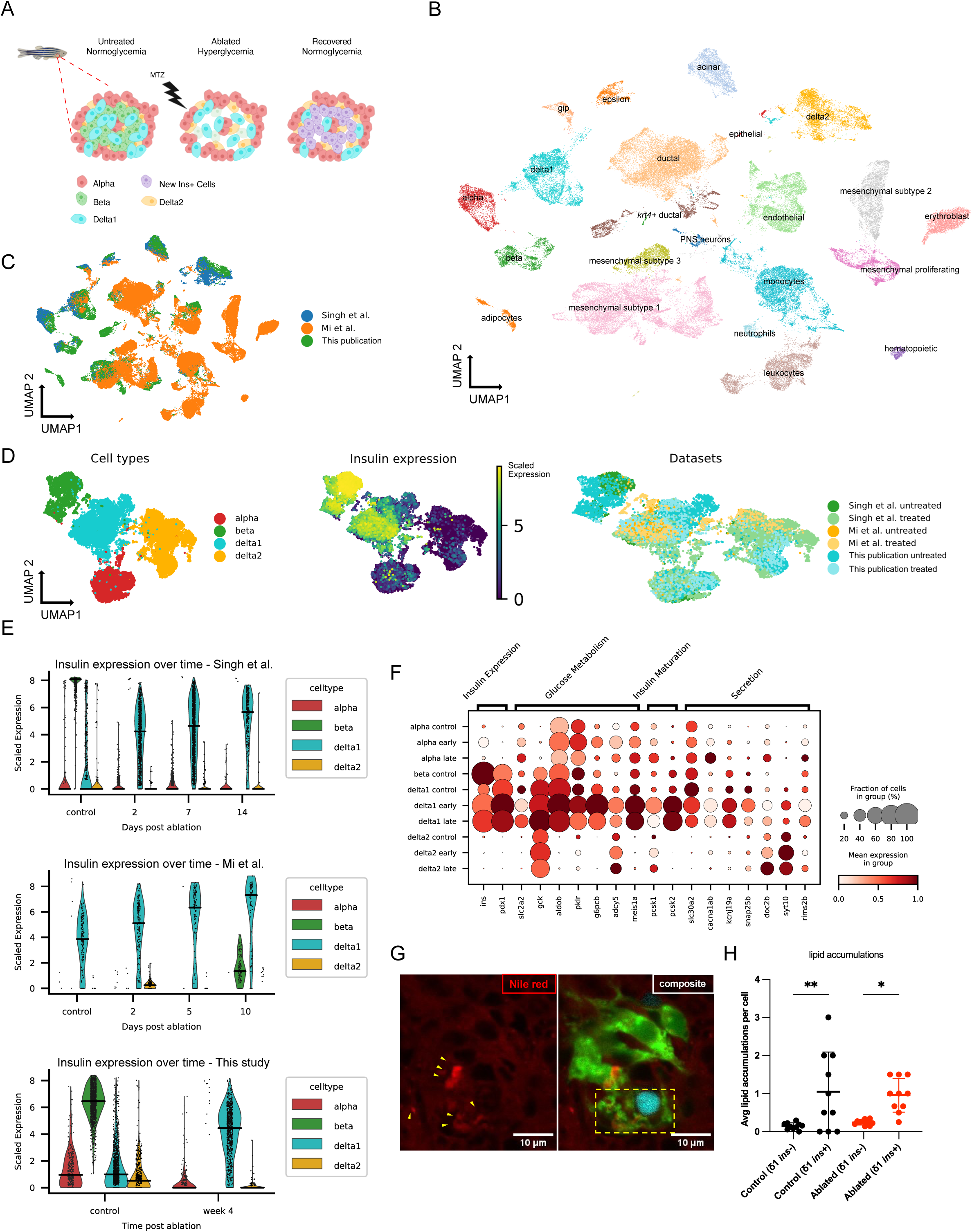
δ1-cells provide early insulin restoration after β-cell loss in adult zebrafish. A) *Ins-NTR* model (Curado *et al*, 2008; Tucker *et al*, 2023) used in studies for targeted β-cell ablation and regeneration in zebrafish. B) UMAP representation of different cell types in scRNA-seq atlas of zebrafish regeneration (n = 85,614 cells). C) Origin of the data: Dataset contains novel data and data from Singh et al. (Singh *et al*, 2022) and Mi et al. (Mi *et al*, 2023). D) Integrated data of treated and untreated endocrine cells (n = 16,426) showing *ins* expression in δ1-cells post β-cell ablation. E) Quantification of *ins* gene expression in endocrine cells post β-cell ablation. Horizontal bar: median. F) Selection of genes relevant for insulin expression, glucose metabolism and hormone secretion differentially expressed in δ1-cells post β-cell ablation. G) Representative confocal image of a larval pancreatic islet after β-cell ablation. δ1-cells are labelled by sst1.1:EGFP-Ras (green), insulin-expressing cells by ins:NLS-mCerulean (cyan), and neutral lipids by Nile Red (red). Yellow arrowheads mark lipid accumulations in δ1-cells; the yellow box outlines a bihormonal ins⁺ δ1-cell. Scale bar, 10 μm. H) Lipid accumulations are abundant in ins^+^ δ1-cells but not in ins^−^ δ1-cells, indicating altered lipid handling during the regenerative response. Each point represents one islet (n = 10 larvae). Statistical significance was determined using one-way ANOVA with Tukey’s multiple comparisons test. Bars indicate mean ±1 SD.

## Results

### A primed δ1-cell state supports early insulin restoration after β-cell loss

To identify the cellular source of restored insulin production after β-cell ablation, we analyzed single-cell RNA-sequencing (scRNA-seq) data from the regenerating adult zebrafish pancreas. While endocrine cells are typically characterized by expression of cell type specific hormones, we reasoned that information about the whole transcriptome would be required to attribute insulin production from unconventional sources to the correct cell types. We integrated two existing scRNA-seq datasets (Singh *et al*, 2022; Mi *et al*, 2023) of adult zebrafish pancreas after β-cell ablation, which we reanalyzed and expanded by adding an additional later time point (1 month post β-cell ablation) (Figure 1B, C). We detected the expected cell types of the endocrine and exocrine pancreas as well as cell populations from adjacent tissues (Figure 1B, S1A). Of note, we observed different proportions of cell types between the three datasets, which can be explained by differences in cell isolation: The dataset by Mi et al. was generated using FACS to isolate *krt4*^+^ lineage traced cells, while no cell selection was applied in Singh et al. and in our newly generated dataset. Cell types present in all datasets integrated well across datasets (Figure 1C).

All datasets contained cells of the endocrine pancreas, specifically β-cells (expressing *ins*), α-cells (*gcga / gcgb*), ε-cells (*ghrl*), and, as previously reported (Singh *et al*, 2022; Carril Pardo *et al*, 2022) two types of δ-cells: δ1-cells expressing *sst1.1*, and δ2-cells expressing *sst2* (Figure 1D). By comparing insulin expression across cell types and time points, we observed that δ1-cells are the main source of insulin post β-cell ablation (Figure 1D). Indeed, we neither observed substantial expression of insulin in any other endocrine cell type, nor reappearance of larger numbers of bona fide β-cells within the first month after β-cell loss in any of the examined datasets (Figure 1E, S1B). Instead, in all datasets a large fraction of the δ1-cell population turned insulin positive upon β-cell ablation (Figure 1E, S2), while maintaining their overall transcriptional identity as δ1-cells (Figure 1D). To further strengthen these observations, we turned to another dataset of bulk RNA-seq of FACS sorted *sst1.1*^+^/*ins*^+^ hybrid cells at 20 dpa (Carril Pardo *et al*, 2022). These cells cluster closely with pseudo-bulked δ1-cells of our integrated dataset (Figure S1C) and show highly increased expression of insulin compared to untreated cells, confirming the δ1 identity of *ins* expressing cells post ablation.

In addition to the increase in insulin, we also observed upregulation of key genes important for glucose metabolism (*slc2a2*, *gck*, *aldob*, *pklr*, *g6pcb, meis1a*), insulin maturation in the form of proprotein convertases (*pcsk1*, *pcsk2*), as well as genes associated with the secretory machinery, like calcium channels (*cacna1ab*, *cacna1da*, *cacna1c*), potassium channels (*abcc8*, *kcnj19a*) and genes related to vesicle trafficking and release (*snap25b*, *doc2b*, *syt1a*, *syt10*, *rims2b*) in δ1-cells upon β-cell ablation (Figure 1F). In line with these observed transcriptional changes related to metabolism, we observed evidence for altered lipid handling in larvae upon β-cell ablation, as suggested by neutral lipid accumulation in insulin-expressing hybrid δ1-cells compared to mono-hormonal *sst1.1*^+^ δ1-cells (Figure 1G,H). Furthermore, inhibition of *ppara*, a central regulator of fatty acid degradation, led to increased formation of bihormonal δ1-cells in ablated as well as unablated conditions (Figure S3), indicative of a functional role of metabolic remodeling.

Despite previous reports that zebrafish regenerate their pancreas by forming novel β-cells (Moss *et al*, 2009; Ghaye *et al*, 2015; Delaspre *et al*, 2015; Mi *et al*, 2023), we did not detect substantial numbers of new β-cells within 30 days after ablation. Instead, early recovery after β-cell loss was mediated predominantly by *sst1.1*^+^ δ1-cells that adopted an insulin-expressing hybrid state while retaining overall δ1-cell identity. Because insulin expression alone would classify these cells as β-like, earlier studies may have misidentified them as newly formed β-cells. Of note, our findings are in line with the previous report that *krt4^+^*-lineage traced cells express insulin after β-cell ablation (Mi *et al*, 2023), as δ1-cells are *krt4*^+^ at 48 hpf (Figure S4). Taken together, these data show that early recovery after β-cell loss is mediated predominantly by *sst1.1*^+^ δ1-cells that adopt an insulin-expressing hybrid state rather than by rapid reconstitution of canonical β-cells.

### A pre-existing accessible chromatin state poises δ1-cells for insulin induction

Because δ1-cells activate insulin expression while retaining overall δ-cell identity, we asked whether they require major epigenetic remodeling or are epigenetically poised to do so. To investigate this hypothesis, we generated scATAC-seq datasets of pancreatic islets at untreated control conditions as well as at 7 and 21 days post β-cell ablation (Methods) in order to compare open chromatin profiles across cell types and time points. We assigned cell types based on gene activity (Methods), and, similar to the analysis of scRNA-seq data in Figure 1, we detected the expected cell types, with a clear separation of the different endocrine cell populations (Figure 2A, S5A). When inspecting chromatin accessibility around transcriptional start sites (TSS) of unablated control samples, we found the canonical endocrine marker genes to be specific for the proposed cell types (Figure 2B). Interestingly, accessibility at the *ins* locus was highest in β-cells, followed by δ1- and α-cells, with δ1-cells showing significantly greater accessibility than α-cells (fold change = 1.3, adjusted p = 1 × 10^−26^). Similarly, when analyzing the gene activity of other β-cell genes that are upregulated in δ1-cells upon β-cell loss (see Figure 1F), we found that δ1-cells displayed more open chromatin compared to α- and δ2-cells, already before β-cell ablation (Figure 2C). Taken together, these findings suggest that δ1-cells possess permissive chromatin and are, at least to some degree, epigenetically primed for insulin expression.

**Figure 2:**
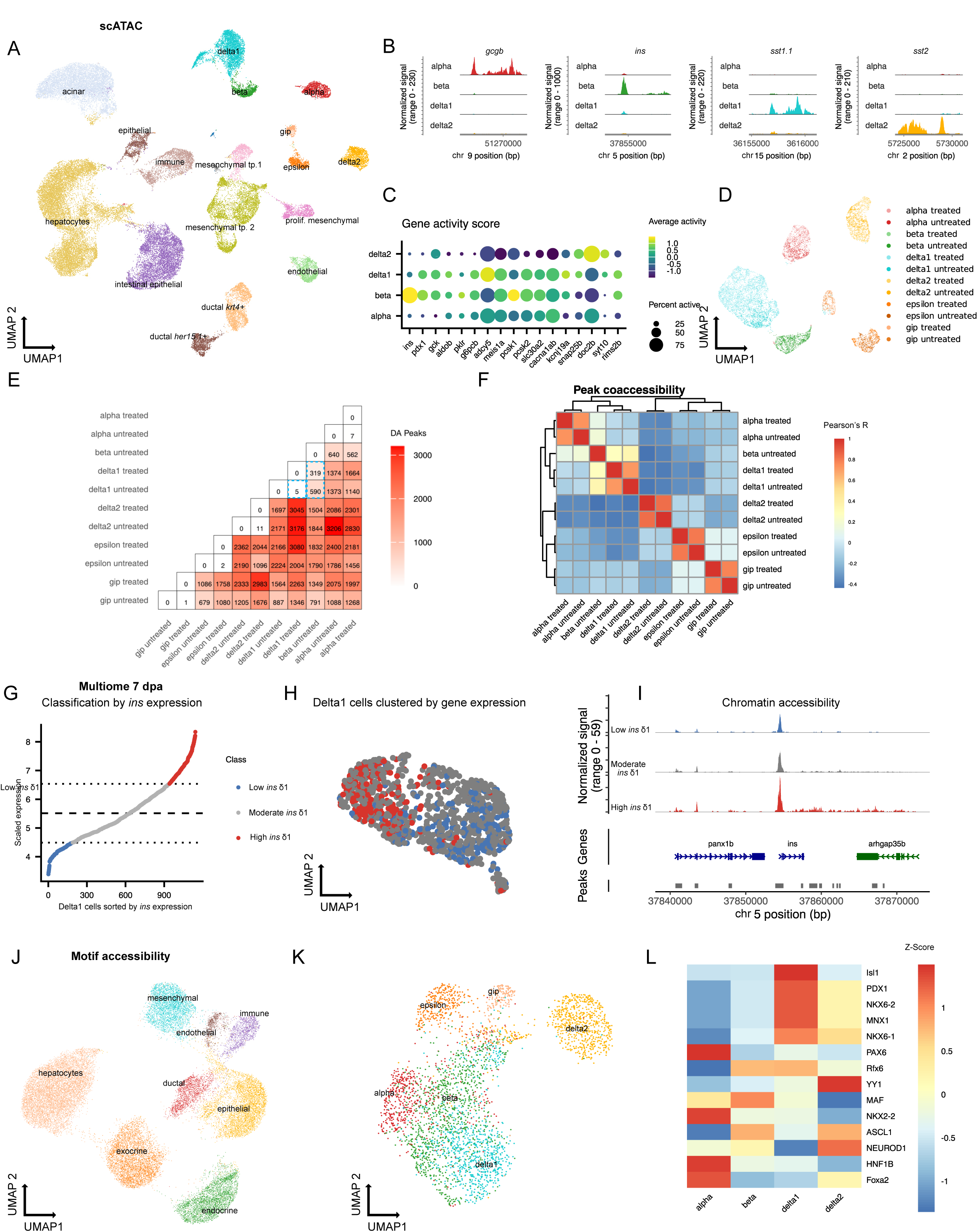
Pre-existing accessible chromatin poises δ1-cells for insulin induction in adult zebrafish. A) UMAP representation of cell types present in the scATAC-seq dataset (n = 51,306 cells). B) Chromatin accessibility around the main endocrine marker genes in untreated control animals. C) Gene activity scores of differentially expressed genes of interest in δ1-cells post β-cell ablation. D) Subset of endocrine cells in the scATAC-seq dataset untreated and post ablation (7 dpa, 21 dpa) (n = 8,819). E) Pairwise comparison of differential peak accessibility in endocrine cells untreated and post ablation (7 dpa, 21 dpa). Logistic regression testing with log_2_ fold change > 1 and Benjamini-Hochberg adjusted p-value of < 0.01. Blue dotted squares highlight the minimal changes between treated and untreated δ1-cells. F) Pearson correlation coefficient of peak co-accessibility between the different cell types. G) Classification of δ1-cells of the multiome data (7 dpa) into *ins_high_* and *ins_low_* using the RNA modality. H) Clustering of δ1-cells of multiome data (7 dpa) by RNA modality allows separation between *ins_high_* and *ins_low_* δ1-cells. I) Comparison of chromatin accessibility at the *ins* gene location between *ins_high_* and *ins_low_* δ1-cells of multiome data (7 dpa). J, K) UMAP representation based on motif accessibility of the different lineages (J) and different endocrine populations (K). L) Z-scores of selected motif accessibilities relevant for β-cell differentiation and function.

We next focused on the chromatin profiles of endocrine cells following β-cell ablation at 7 and 21 dpa (Figure 2D). As expected, β-cells were absent at both timepoints. We determined differential peaks for each pairwise combination of endocrine cell types pre- and post β-cell ablation in order to quantify similarity of open chromatin profiles across cell types and conditions (Figure 2E, S5B). This analysis revealed that there are no significant changes in chromatin accessibility in δ1-cells before versus after ablation, suggesting that insulin induction does not require major chromatin remodeling. Similarly, we also did not observe major chromatin remodeling in any other endocrine cell type post β-cell ablation. Comparing the different endocrine cell types, we found that δ1-cells were most similar to β-cells, exhibiting fewer differential peaks (280) than δ2- (1780) and α- (514) cells. Correlation analysis of open chromatin profiles (Figure 2F) further revealed high similarity between β-cells and δ1- and α-cells, but not δ2, ε, or gip cells (*glucose-dependent insulinotropic peptide responsive cells)*, underscoring the close similarities between δ1- and β-cells.

To further investigate δ1-cell identity, we performed paired scATAC/scRNA-seq multiome sequencing, enabling us to compare chromatin accessibility between *ins*⁺/*sst1.1*⁺ and *ins^−^* /*sst1.1*⁺ δ1-cells at 7 dpa. While the scATAC modality alone did not allow us to differentiate between *ins*⁺ and *ins^−^* δ1-cells (Figure S6), this was possible using the scRNA modality (Figure 2G,H). *Ins*⁺/*sst1.1*⁺ δ1-cells show some opening of the chromatin at the *ins* locus (Figure 2I), but no significant differential peak accessibility compared to *ins*⁻*/sst1.1*⁺ δ1-cells. These data indicate that δ1 competence reflects a pre-existing permissive chromatin state rather than injury-induced large-scale chromatin remodeling.

To better understand the regulatory landscape of δ1-cells, we compared differences in motif accessibility between single cells (Methods). We found that endocrine cells have closely related motif enrichment signatures, but are clearly distinguished from other lineages in the dataset (Figure 2J). Focusing on the endocrine pancreas, we observed that motif enrichment allowed differentiating between the different cell types (Figure 2K). When comparing β-cell related motif accessibility, we discovered high similarity between δ1- and β-cells (Figure 2L, S5C/D). Motifs associated with key endocrine and β-cell regulators, including *isl1*, *nkx6*.*1*, *mnx1* and *pdx1* (Prince *et al*, 2017), were already highly accessible in untreated δ1-cells, further supporting the close regulatory similarity between δ1- and β-cells. Together, these data indicate that δ1-cells are not induced to become permissive after injury, but instead occupy a pre-existing accessible state that licenses rapid endocrine plasticity.

### Cross-species comparison separates permissive, deployable and blocked endocrine states

To distinguish regulatory features associated with regenerative competence from those that are more broadly conserved, we compared endocrine chromatin states across zebrafish, medaka and human pancreas. We first focused on medaka (*Oryzias latipes*), another teleost model in which β-cell regeneration has been examined (Otsuka & Takeda, 2017). We collected scATAC-seq data of the Medaka pancreas (Figure 3A, Figure S7) and annotated the cells based on known markers – *gcgb* for α-cells, *ins* for β-cells, *sst1.1* for δ1-cells and *sst1.2* for δ2-cells (Figure 3B, Figure S7D). Medaka has two paralogous insulin genes, *ins1.1* and *ins1.2*. Interestingly, we observed open chromatin at the *ins1.2* gene in medaka δ1-cells, similar to the permissive open chromatin signature in zebrafish. By contrast, open chromatin for *ins1.1* was exclusively detected in β-cells (Figure 3C).

**Figure 3:**
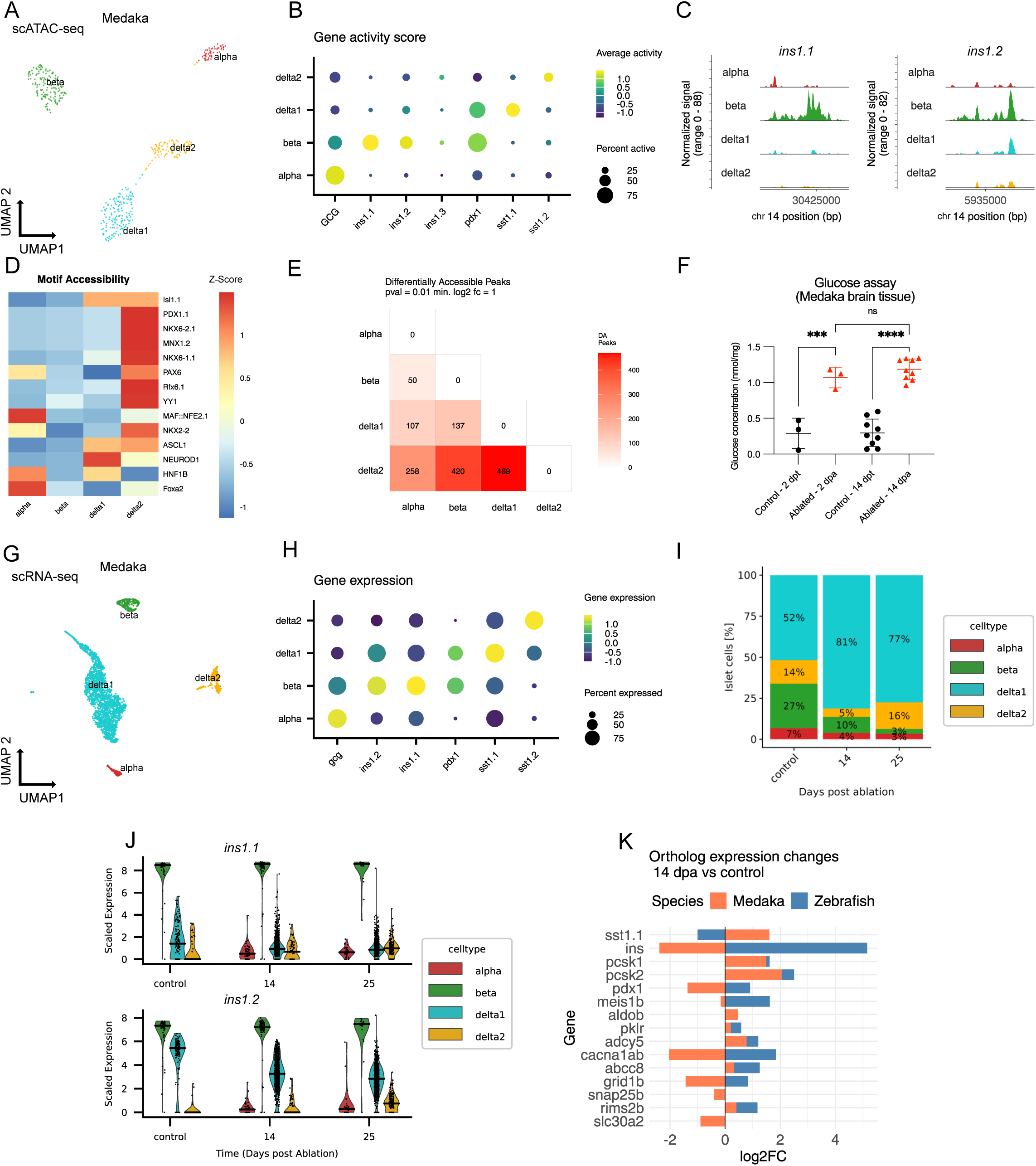
Partially permissive chromatin state but absence of hybrid δ1-cells in the medaka pancreas post β-cell ablation. Single-cell chromatin accessibility (A–E) and transcriptomic (G–K) profiling of the medaka pancreatic islet. A) UMAP representation of scATAC-seq data of medaka islets. B) Chromatin accessibility in medaka around the main endocrine marker genes. C) Chromatin accessibility of the two active insulin orthologues in medaka. D) Z-scores of selected motif accessibilities relevant for β-cell differentiation and function in medaka. E) Pairwise comparison of differential peak accessibility in endocrine cells of the unablated medaka pancreas. Logistic regression testing with log_2_ fold change > 1 and Benjamini-Hochberg adjusted p-value of < 0.01. F) Blood glucose levels in adult medaka remain elevated after β-cell ablation at 14 dpa. Control: treated with DMSO. Dpt: Days post treatment. G) UMAP of medaka islet scRNA-seq data from control, 14 dpa, and 25 dpa samples. H) Gene expression of endocrine marker genes in medaka islets. I) Proportion of each islet cell type in control, 14 dpa, and 25 dpa samples. J) Expression of the two active insulin genes in islet cells at control, 14 and 25 days post ablation K) Log2 fold change of genes involved in insulin secretion (as in Fig. 1F) in δ1-cells before and after β-cell ablation, compared between medaka and zebrafish at 14 dpa.

We next compared motif accessibility of key transcription factors relevant for endocrine differentiation. Motif analysis revealed both shared endocrine regulatory features and clear differences between δ1- and δ2-cells, as well as divergence between corresponding endocrine populations in zebrafish and medaka (Figure 3D, Figure S7F). Quantification of differentially accessible peaks revealed that δ1-cells are closer to β-cells than to δ2-cells (Figure 3E). Taken together, in medaka we find evidence for permissive chromatin, similar to the zebrafish. This finding motivated us to perform a functional experiment in which we performed NTR ablation of β-cells in adult medaka. Based on measurement of glucose concentration, we did not find evidence for functional recovery of insulin production at 14 dpa (Figure 3F). We next investigated the transcriptional response of medaka δ1-cells to β-cell ablation. scRNA-seq allowed identification of the endocrine cell types at 14 and 25 dpa (Figure 3G, H, S8), including a strong reduction of β-cells after MTZ treatment (Figure 3I). In line with the observed lack of functional regeneration, we did not detect insulin production in medaka δ1-cells at 14 and 25 dpa (Figure 3J). Scoring of genes that are upregulated in zebrafish δ1-cells upon β-cell ablation did not show a consistent trend in the medaka dataset (Figure 3K), underscoring the differential response to β-cell loss. Thus, medaka δ1-cells show a permissive chromatin state similar to zebrafish for *ins1.2*, but this state is insufficient on its own to enable adult functional recovery.

Cellular plasticity of adult human endocrine cells is believed to be very limited, and the existence of insulin-producing hybrid δ cells has not been reported. We next sought to understand to which degree this could be explained by a lack of permissive chromatin. To this end, we gathered available paired scRNA/scATAC-seq data from non-diabetic donors collected in the *human pancreas analysis program* (Patil *et al*, 2023) (Figure 4A, S9) and annotated the cells based on the scRNA-seq modality (Figure 4B). In agreement with previous reports, we found a single population of δ cells in the human pancreas (Segerstolpe *et al*, 2016), with motif accessibility closely matching zebrafish δ1-cells (Figure 4C), which indicates a high degree of evolutionary conservation. Similar to zebrafish and medaka, differential peak accessibility revealed that human δ-cells are relatively close to β-cells on the level of open chromatin (Figure 4D). However, in contrast to zebrafish and medaka, the accessibility of the *INS* locus is highly restricted in human δ-cells (Figure 4E, F). The restricted accessibility of the *INS* locus in human δ-cells points to an epigenetic barrier that may limit endocrine plasticity in the adult human pancreas. Together, the zebrafish–medaka–human comparison defines three regulatory states of endocrine plasticity: a permissive and deployable state in zebrafish, a permissive but insufficient state in medaka, and a restricted state in human δ-cells. This comparative framework suggests that regenerative competence is determined not simply by lineage proximity, but by whether permissive chromatin can be productively deployed after injury.

**Figure 4:**
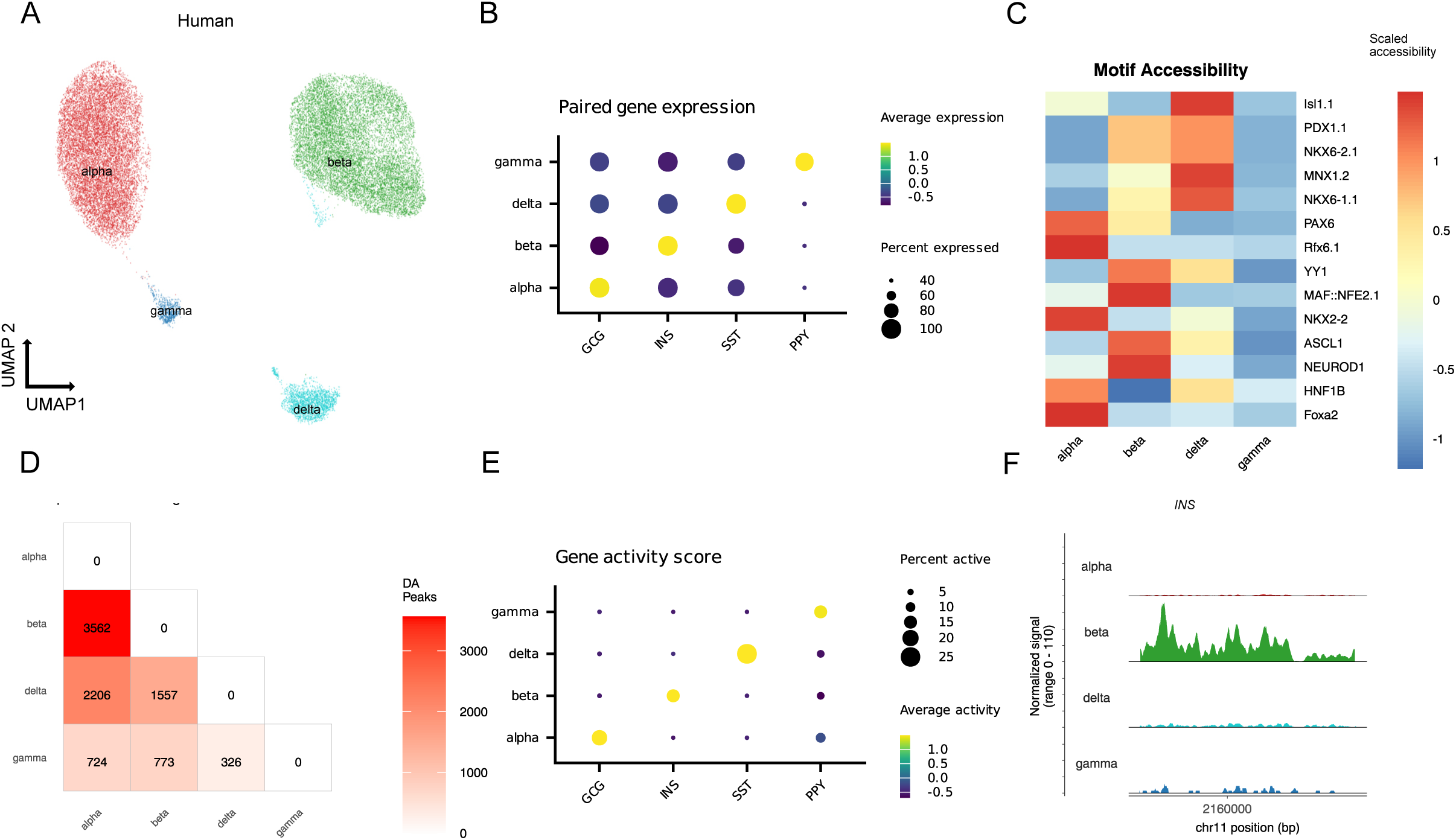
Restrictive chromatin state in the human pancreas. A) UMAP representation of scRNA-seq/scATAC-seq multiome sequencing data of non-diabetic human islets. B) Paired expression of selected marker genes in human islets. C) Z-scores of selected motif accessibilities relevant for β-cell differentiation and function in human islets. D) Pairwise comparison of differential peak accessibility in endocrine cells of non-diabetic human islets. Logistic regression testing with log_2_ fold change > 1 and Benjamini-Hochberg adjusted p-value of < 0.01. E) Chromatin accessibility around the main endocrine marker genes in human islets. F) Chromatin accessibility of the *INS* gene in non-diabetic adult human.

### Gene regulatory network analysis identifies instructive regulators of δ1 state switching

Because permissive chromatin alone could not explain injury-specific insulin induction in δ1-cells in the case of medaka, we next searched for instructive transcription factors that might drive this response. We integrated pseudobulk scRNA-seq and scATAC-seq data to link differentially expressed genes to candidate cis-regulatory elements (cCRE) and then used motif enrichment and TF-expression correlations to infer candidate TF–cCRE–gene relationships (Figure 5A, S10; Methods). This analysis identified transcription factors whose predicted regulatory activity increased in δ1-cells after β-cell ablation. We focused our analysis on genes differentially expressed in endocrine cells after the β-cell ablation in different timepoints (Figure 5B). As expected, the linked cCREs were mostly also accessible in untreated δ1-cells (Figure 5C). Using the TF-cCRE pairs, we then calculated the enrichment of putative TF binding sites at the different timepoints tested (Figure 5D, S11). This analysis prioritized *pdx1*, *meis1a*, *meis1b*, *prdm1b* and *pbx3b* as candidate regulators of the δ1 gene expression switch. Among these, *pdx1* served as an expected positive control given its established role in β-cell differentiation and insulin expression (Prince *et al*, 2017). By contrast, *meis1a/b* stood out as strong and unexpected candidates because *MEIS1* has been linked to metabolic state control in other systems, including promotion of glycolytic programs and suppression of oxidative phosphorylation (Simsek *et al*, 2010; Kocabas *et al*, 2012). Consistent with this, many genes induced in δ1-cells after β-cell loss are metabolism-related (Figure 1F), *meis1a*-linked cCREs became increasingly accessible after ablation (Figure 5E), and predicted *meis1a* target genes were enriched for glycolysis and gluconeogenesis (Figure 5F, S12). In contrast to zebrafish, we did not observe upregulation of meis1a/b in medaka δ1-cells upon β-cell loss at 14 and 25 dpa (Figure 3K). Furthermore, in pancreatic development, Meis proteins are expressed in the endocrine pancreas and cooperate with Pbx factors to regulate Pax6 enhancer activity (Zhang *et al*, 2006). These analyses nominate *meis1a*/*b* as candidate instructive regulators that link metabolic remodeling to insulin induction in δ1-cells.

**Figure 5:**
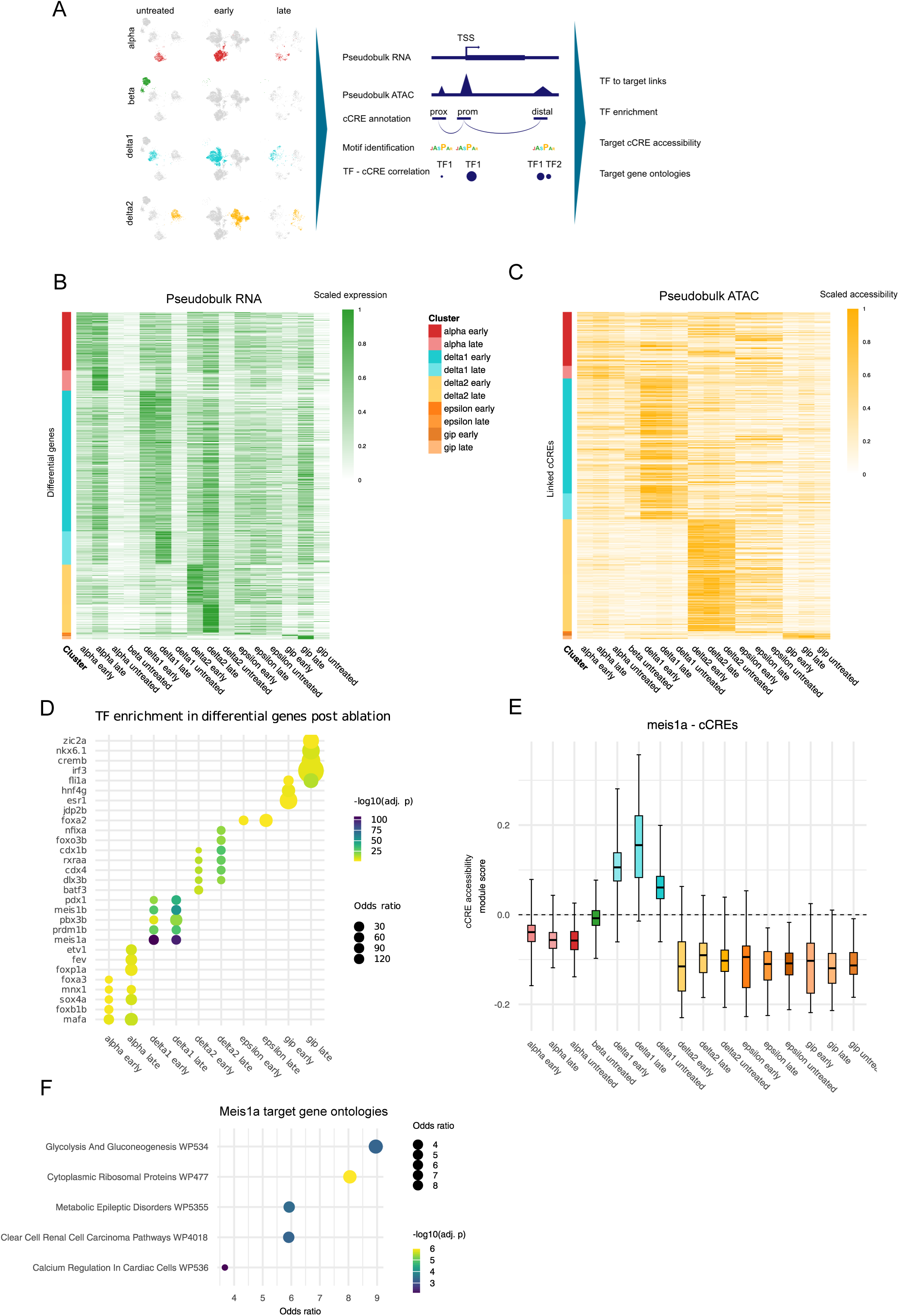
*meis1a/b* emerges as a metabolic regulator of δ1-cell plasticity in zebrafish. A) Schematic of the GRN construction. B) Heatmap of pseudobulk RNA data of genes differentially expressed post β-cell ablation. C) Linked pseudobulk cCRE accessibility of elements linked to differential gene expression post ablation. D) Enrichment of TF-Gene links in gene sets differentially expressed post ablation. E) Module accessibility of cCREs linked to *meis1a* in different cell types and conditions. F) Gene ontologies of differentially expressed genes linked to the transcription factor *meis1a*.

### meis1a/b is required for deployment of the hybrid endocrine program

To functionally test candidate regulators identified by our gene regulatory network (GRN) analysis, we next performed targeted perturbation experiments in zebrafish larvae. We used the larval β-cell ablation model as a rapid in vivo assay to test whether candidate regulators identified in the adult regenerative response are required for hybrid-cell formation. From the set of transcription factors predicted to regulate genes induced in δ1-cells after β-cell ablation – including *pdx1*, *meis1a*, *meis1b*, *prdm1b*, and *pbx3b* – we prioritized candidates that showed strong motif enrichment in regulatory elements linked to differentially expressed genes and that were expressed in δ1-cells during the regenerative response (Figure 5D–F).

We generated CRISPR–Cas9 crispants (Kroll *et al*, 2021), targeting selected candidate transcription factors and quantified the formation of insulin-expressing hybrid cells following β-cell ablation. Larvae were subjected to nitroreductase-mediated β-cell ablation and analyzed during the early phase of recovery, when δ1-derived hybrid cells normally emerge (Figure 6A).

**Figure 6:**
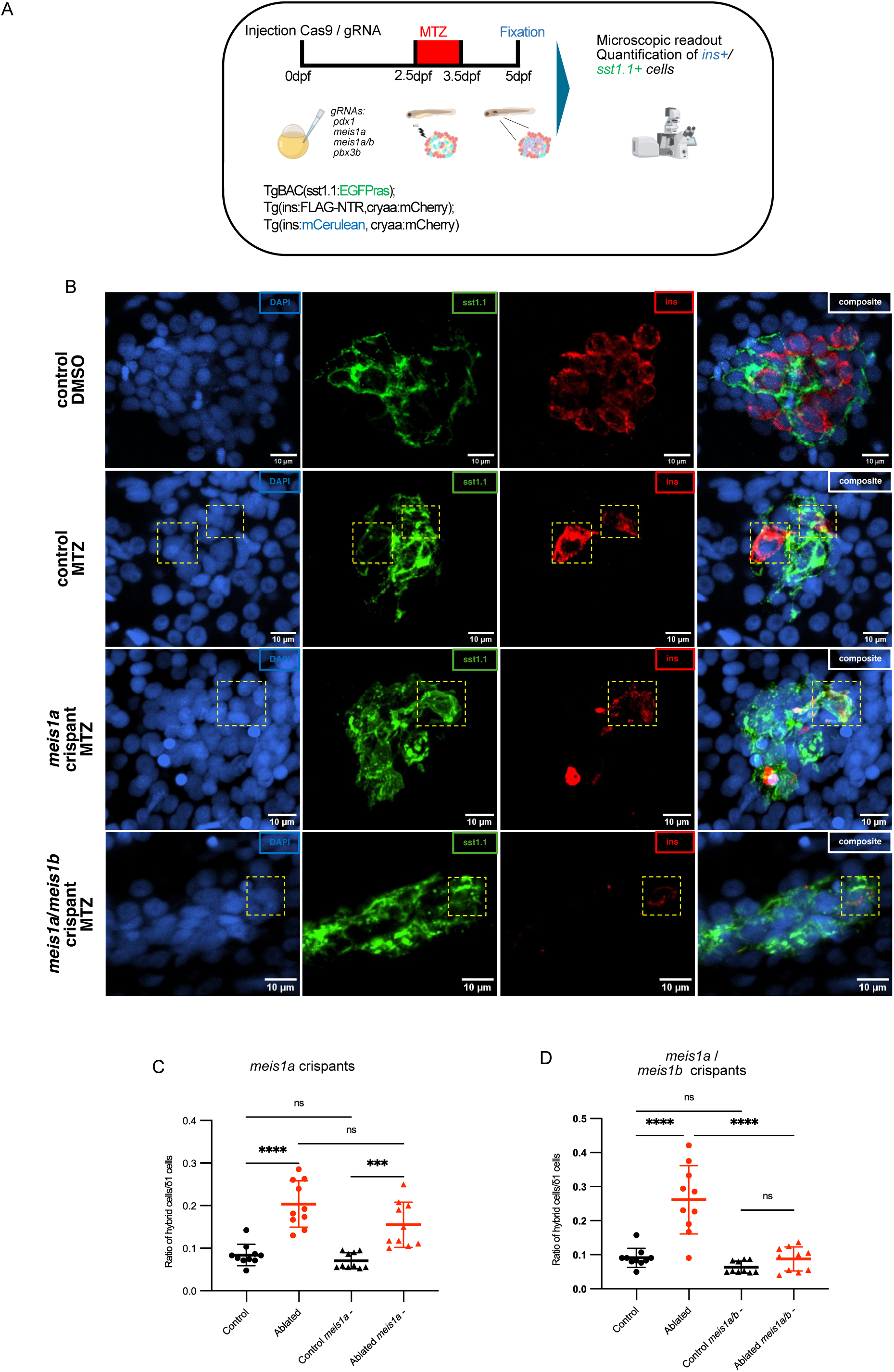
*meis1a/b* is required for injury-induced insulin induction in δ1-cells in zebrafish larvae. A) Experimental strategy to test candidate transcription factors regulating δ1 hybrid cell formation. Candidate regulators identified by gene regulatory network analysis (Figure 5) were perturbed using CRISPR–Cas9–mediated genome editing in zebrafish larvae. β-cells were ablated using the insulin promoter–driven nitroreductase (Ins:NTR) system, and the formation of insulin-producing hybrid cells was quantified during the early recovery phase. B) Representative confocal images of pancreatic islets pre and post β-cell ablation, and after F0 knockout of *meis1a* and *meis1a/meis1b*. In MTZ ablated control larvae, δ1-cells (*sst1.1*^+^) activate insulin expression and form bihormonal hybrid cells (yellow dotted squares). F0 knockout of *meis1a* alone does not significantly reduce the number of insulin-expressing δ1-cells. In contrast, combined F0 knockout of *meis1a* and *meis1b* markedly reduces the number of insulin-expressing δ1-cells. Scale bar, 10 µm. C) Quantification of δ1 hybrid cell formation following candidate TF perturbation. CRISPR perturbation of meis1a does not significantly reduce the number of *sst1.1^+^/ins^+^* hybrid cells compared with controls. Each point represents one islet (n = 10 larvae). Statistical significance was determined using one-way ANOVA with Tukey’s multiple comparisons test. D) CRISPR perturbation of both *meis1a* and *meis1b* significantly reduces the number of *sst1.1^+^/ins^+^* hybrid cells compared with controls, identifying *meis1a/meis1b* as required regulators of insulin induction in δ1-cells following β-cell loss.

As expected, due to its role as a key factor in the development of the islet (Prince *et al*, 2017), *pdx1* F0 crispants showed large disruptions in the islets (Figure S13), confirming the efficiency of our knockout model. Perturbation of *meis1a* as well as combined perturbation of both *meis1a*/*b* did not disrupt islet development, and both δ1- and β cells were present in the islets (Figure 6B). While *meis1a* perturbation alone did not significantly reduce *ins* expression by δ1-cells post β-cell ablation (Figure 6C), combined *meis1a/b* perturbation led to a pronounced reduction in insulin production. While control larvae robustly formed bihormonal *sst1.1*^+^/*ins*^+^ cells following β-cell ablation, *meis1a/b* crispants displayed a significant reduction in hybrid cell formation relative to controls (Figure 6D), indicating that *meis1a*/*b* activity is required for the activation of insulin expression in δ1-cells during pancreatic recovery. Importantly, perturbation of *meis1a*/*b* did not abolish the presence of δ1-cells themselves, suggesting that *meis1a*/*b* specifically affects the transcriptional transition toward the hybrid endocrine state rather than δ1-cell survival or identity. Together, these findings identify *meis1a/b* as instructive regulators of insulin induction in δ1-cells following β-cell loss.

## Discussion

Our data identify a mechanism by which differentiated adult endocrine cells acquire plasticity after injury. Rather than requiring large-scale rebuilding of cell identity, early insulin restoration after β-cell loss proceeds through deployment of a pre-existing permissive chromatin state and *meis1a/b*-dependent transcriptional control within δ1-cells. In this framework, zebrafish δ1-cells represent a deployable plastic state that is absent or constrained in other vertebrate systems.

Compared with mammalian models of extreme β-cell loss, in which insulin production by non-β endocrine cells is generally limited and gradual, the zebrafish response is strikingly rapid and functionally effective, and is mediated predominantly by δ1-cells that adopt a stable hybrid state rather than undergoing immediate full β-cell conversion (Thorel *et al*, 2010; Cigliola *et al*, 2018). Independent studies have further identified ductal Ngn3-expressing progenitors as contributors to adult β-cell neogenesis in diabetic mice (Gribben *et al*, 2021), indicating that multiple cellular routes to β-cell recovery may coexist depending on injury context. In zebrafish, earlier work likewise supported contributions from ductal compartments (Delaspre *et al*, 2015; Mi *et al*, 2023). Our data do not exclude such contributions in parallel or at later stages, but indicate that within the first month after adult β-cell ablation the dominant insulin-expressing population retains δ1 identity. Specifically, it is possible that recently differentiated δ1-cells from the duct adopt a bi-hormonal state, in addition to pre-existing δ1-cells that start to express insulin. This is in line with previous analysis of insulin and somatostatin protein expression, showing that the bi-hormonal cells persist long-term, until at least 14 dpa (Singh *et al*, 2022) and up to 120 dpa (Carril Pardo *et al*, 2022), indicating that they represent a stable cellular state. In this light, krt4-based lineage tracing (Mi *et al*, 2023) should be interpreted cautiously, because krt4 marks a broader responsive lineage that includes δ1-cells. Whether the previously described Notch-responsive population (Delaspre *et al*, 2015) is molecularly distinct from, overlaps with, or gives rise to the primed δ1 state identified here remains an open question. In medaka we did not observe formation of abundant hybrid cells post β-cell ablation. Instead, our scRNA seq only detected a reduced population of mono hormonal insulin-expressing β-cells at 14 and 25 dpa. This population may represent cells that slowly differentiate from the duct, or are spared β-cells that survived the ablation. In contrast to zebrafish, medaka remained hyperglycemic, suggesting that the lack of abundant hybrid cell formation hinders effective restoration of insulin production and the return to normoglycemia.

A central conceptual advance of our study is the separation of permissive and instructive features underlying this response. On the permissive side, δ1-cells are epigenetically unusually close to β-cells: they show relatively open chromatin at *ins* and other β-cell-associated loci even before injury, consistent with a poised state for insulin activation. β-cell loss does not trigger large-scale remodeling of accessibility, implying that the switch into a hybrid state is driven more by deployment of pre-existing regulatory potential than by major reconfiguration of chromatin. This does not exclude important regulation below the resolution of accessibility maps, including altered transcription factor occupancy, enhancer-promoter communication, or histone modifications, but it suggests that rapid insulin induction does not require the de novo opening of a previously inaccessible endocrine program.

This framework also helps explain the interspecies comparison. Medaka δ1-cells appear to share a permissive chromatin configuration at insulin loci, yet adult medaka failed to recover normoglycemia in the time window examined, indicating that permissive chromatin is not sufficient on its own. By contrast, human δ-cells lack accessible chromatin at the *INS* locus despite broader regulatory similarities to teleost δ1-cells, suggesting an additional epigenetic barrier to endocrine plasticity in the adult human pancreas. Together, these observations argue that regenerative competence depends on both a permissive chromatin state and injury-responsive instructive inputs.

Our gene-regulatory network analysis nominates such instructive inputs. Alongside expected endocrine regulators such as *pdx1*, we identified *meis1a/b*, *prdm1b* and *pbx3b* as candidate drivers of the δ1 response, and larval perturbation experiments functionally prioritized *meis1a/b* as required for insulin production by δ1-cells. This result is notable because it places a TALE homeobox factor not simply downstream of endocrine identity change, but near the top of the regulatory hierarchy that enables the hybrid state.

MEIS1 is especially compelling in light of prior work linking it to metabolic state control in other systems. Long-term hematopoietic stem cells rely on glycolysis rather than mitochondrial oxidative phosphorylation, and Meis1 contributes to this state through transcriptional control of Hif-1α; subsequent work showed that Meis1 also supports Hif-1α/Hif-2α-dependent oxidant defense, with Meis1 loss shifting HSCs toward mitochondrial oxidative metabolism and loss of quiescence (Simsek *et al*, 2010; Kocabas *et al*, 2012). In a second context, normal down-regulation of MEIS1 during the perinatal period promotes maturation of oxidative phosphorylation in cardiomyocytes (Lindgren *et al*, 2019). Finally, in pancreatic cancer cells, Meis1 has been reported to directly regulate mitochondrial gene transcription via binding to the mitochondrial H-strand promoter (Tomoeda *et al*, 2011). Although these systems are biologically distinct, they converge on a common theme: MEIS1 can couple cell-state transitions to control of glycolysis, oxidative metabolism and mitochondrial gene expression.

That theme fits remarkably well with our data. Following β-cell ablation, δ1-cells upregulate not only insulin, but also genes involved in glucose uptake and metabolism, prohormone processing, ion handling and vesicle release, and they accumulate lipids. Moreover, interference with *Ppara*-driven fatty-acid metabolism enhances hybrid-cell formation, arguing that metabolic state is not merely a consequence of the fate change but may be part of the mechanism enabling it. In this context, *meis1a*/*b* may provide a link between injury-induced metabolic stress and transcriptional activation of a β-like program. The enrichment of glycolysis/gluconeogenesis-related genes among predicted *meis1a* targets is consistent with such a model. We therefore favor a view in which *meis1a*/*b* do not simply switch on insulin transcription, but help reconfigure δ1-cells into a metabolic state compatible with insulin synthesis and secretion after β-cell loss.

This interpretation also sharpens the meaning of regeneration in this system. In the time frame examined, the zebrafish pancreas does not rapidly recreate its pre-injury cellular composition; instead, it restores function through a stable or semi-stable hybrid state. An important next question is whether bona fide β-cells subsequently re-emerge during longer recovery.

More broadly, our study suggests that successful endocrine plasticity requires alignment of three regulatory layers: a permissive chromatin landscape, injury-responsive transcription factors, and potentially a metabolic state capable of communicating β-cell loss. In zebrafish, δ1-cells satisfy all three conditions; in medaka, permissive chromatin appears present but may lack the appropriate instructive circuitry; in humans, the *INS* locus itself appears epigenetically restricted. For translational efforts, this argues against strategies focused only on lineage transcription factors or only on chromatin opening. A more plausible route may be to combine chromatin unlocking with manipulation of metabolic and transcriptional regulators, including *MEIS1*-linked pathways. Whether such interventions can safely induce an insulin-competent state in mammalian δ-cells remains open, but the present work provides a mechanistic framework for asking that question.

## Acknowledgements

We acknowledge the MDC/BIMSB bioinformatics platform, the CMCB and CRTD core facilities: (flow cytometry, light microscopy, histology, and zebrafish). We are grateful to Dr. Christopher Wright for sharing the anti-Pdx1 antibody and the Dresden Concept Genome Center for close collaboration on the project.

## Author Contributions Statement

Conceptualization and study design: J.P.J., N.N., A.F., P.C. and J.G.; Investigation: J.G., P.C., P.S., A.R., E.W.; Formal analysis: J.G., P.S.; Study supervision: J.P.J., N.N., A.F.; Writing – Original draft: J.P.J. and J.G. in close interaction with N.N.; Writing – Editing and Review: all authors.

## Materials and Methods

### Software and algorithms

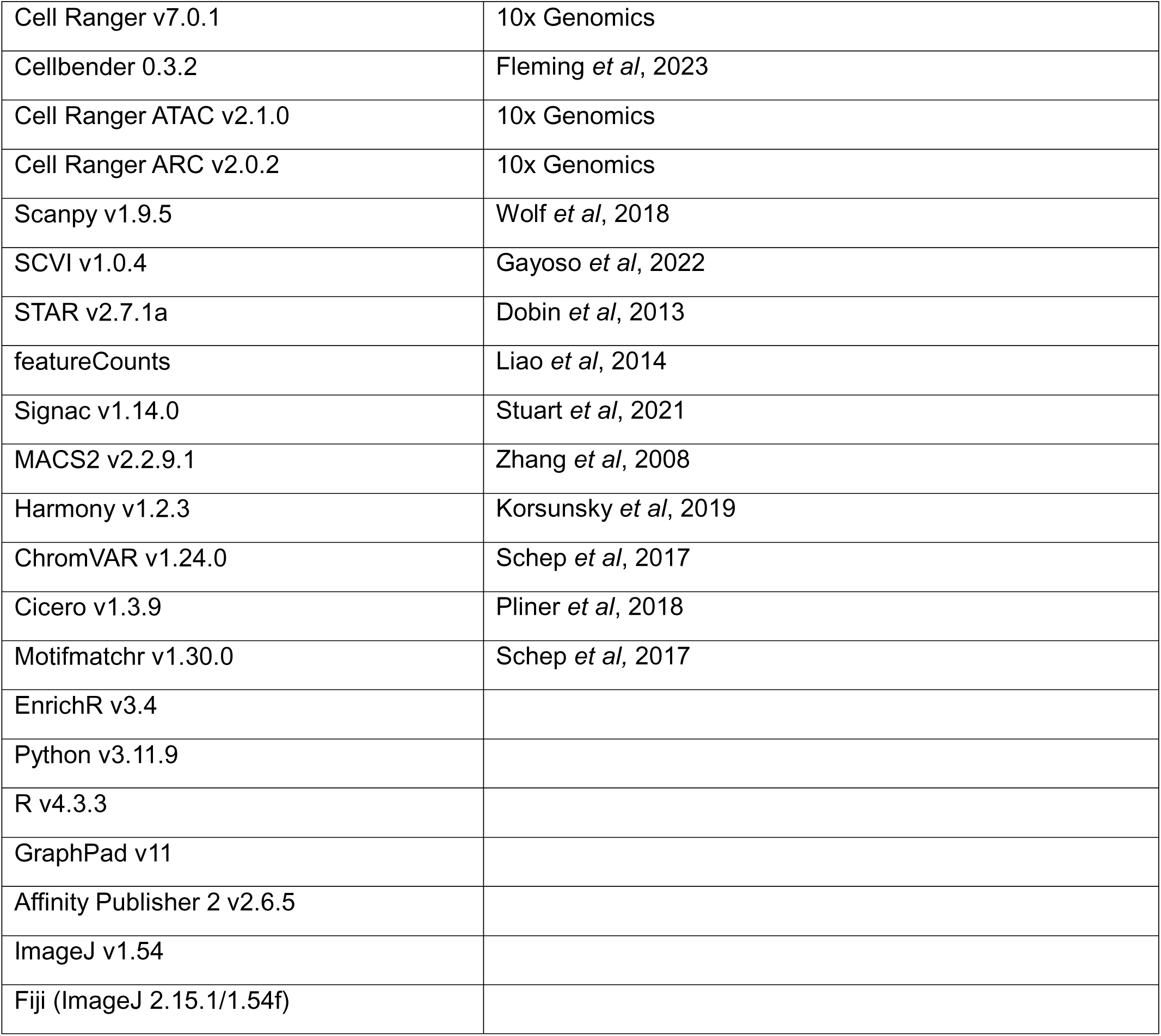

|  |  |
| --- | --- |
| Cell Ranger v7.0.1 | 10x Genomics |
| Cellbender 0.3.2 | Fleming *et al*, 2023 |
| Cell Ranger ATAC v2.1.0 | 10x Genomics |
| Cell Ranger ARC v2.0.2 | 10x Genomics |
| Scanpy v1.9.5 | Wolf *et al*, 2018 |
| SCVI v1.0.4 | Gayoso *et al*, 2022 |
| STAR v2.7.1a | Dobin *et al*, 2013 |
| featureCounts | Liao *et al*, 2014 |
| Signac v1.14.0 | Stuart *et al*, 2021 |
| MACS2 v2.2.9.1 | Zhang *et al*, 2008 |
| Harmony v1.2.3 | Korsunsky *et al*, 2019 |
| ChromVAR v1.24.0 | Schep *et al*, 2017 |
| Cicero v1.3.9 | Pliner *et al*, 2018 |
| Motifmatchr v1.30.0 | Schep *et al*, 2017 |
| EnrichR v3.4 |  |
| Python v3.11.9 |  |
| R v4.3.3 |  |
| GraphPad v11 |  |
| Affinity Publisher 2 v2.6.5 |  |
| ImageJ v1.54 |  |
| Fiji (ImageJ 2.15.1/1.54f) |  |

### Fish strains and husbandry

All experiments were performed in wild-type or transgenic zebrafish of the outbred AB, WIK, or hybrid WIK/AB backgrounds. Fish were maintained under standard conditions at 28°C and assigned randomly to experimental groups. Published transgenic strains used included Tg(ins:BB1.0L; cryaa:RFP) (Singh *et al*, 2017), Tg(sst1.1:EGFP-Ras)^fr40Tg^ (Löhr *et al*, 2018), Tg(ins: FLAG-NTR)^s950^ (Andersson *et al*, 2012), Tg(ins:YFP-2A-NTR3,cryaa: mCherry)^tud201Tg^, Tg(ins:NLS-mCerulean, cryaa:mCherry) (Singh *et al*, 2017).

Medaka were maintained under standard conditions at 26°C and assigned randomly to experimental groups. The non-transgenic medaka strain d-rR.YHNI (Scholz *et al*, 2003) was used in the scRNAseq experiments, hemizygous transgenic animals of TG1679 (Otsuka & Takeda, 2017) carrying the insulin:EGFP-NTR sequence were used for the ablation experiments.

All experiments adhered to the Animal Welfare Act and were approved by the Landesdirektion Sachsen, Germany. (TVV32_2020, TVT1_2021, TVV62_2024).

### Construction of Tg(ins:NLS-mRuby-T2A-NTR2.0, myl7:NLS-mCherry) plasmid

mRuby3 was amplified from addgene plasmid 112169 using Fw-EcoRI-NLS-mRuby and Rev-SpeI-T2A-mRuby primers. The NTR2.0 gene was amplified with the Fw-SpeI-NTR2.0 and Rv-PacI-NTR2.0 primers. The plasmid ins-sepHluorin-cmcl2-XbaNLSmCherry with I-SceI meganuclease sites was used as the backbone and digested with EcoRI and PacI.

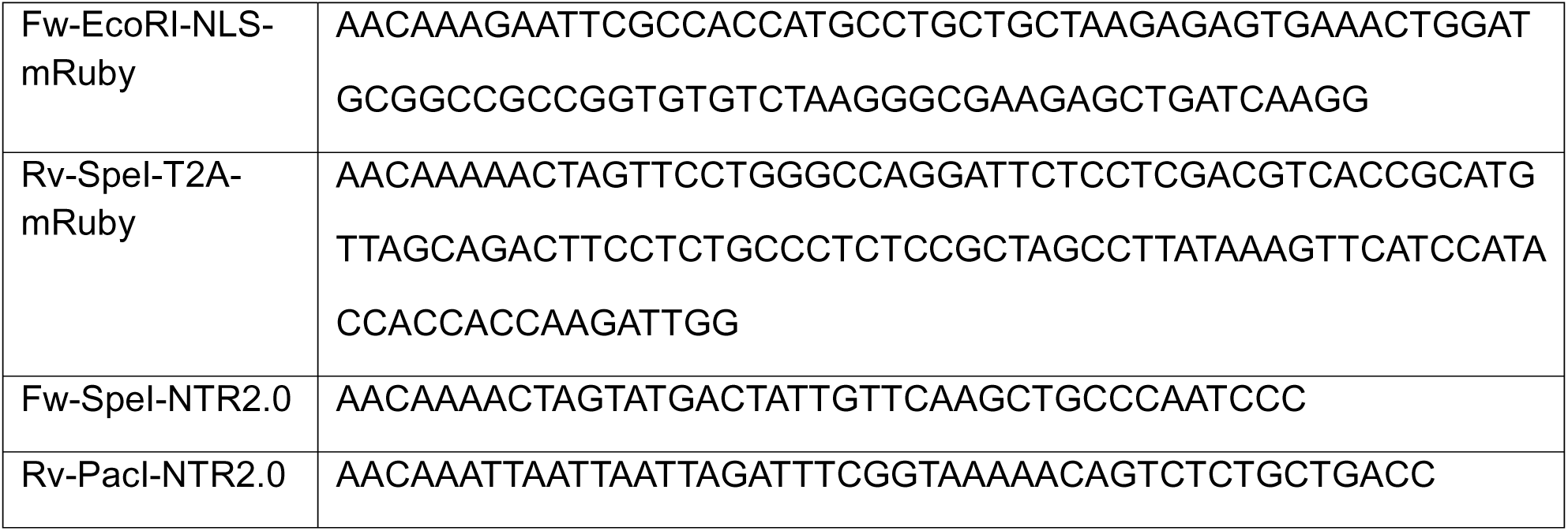

Transgenic founders were generated using the I-SceI meganuclease.

### Crispant F0 knockout perturbation experiments

F0 TF perturbations were performed as previously described (Kroll *et al*, 2021) on TgBAC(sst1.1:EGFPras); Tg(ins:NLS-mRuby-T2A-NTR2.0,myl7:NLS-mCherry) embryos, with the modification that two loci per gene were targeted. crRNAs were designed to target the specific exons listed below using the Alt-R Custom Cas9 Design Tool. The crRNA constituted the only locus-specific component of the Cas9/gRNA ribonucleoprotein (RNP). crRNAs were selected based on a balance of highest predicted on-target activity and minimal off-target potential. Synthetic gRNAs were assembled from separate crRNA (Alt-R CRISPR-Cas9 crRNA) and tracrRNA (Alt-R CRISPR-Cas9 tracrRNA, IDT, Cat. #1072532) components. crRNA/tracrRNA annealing and gRNA/Cas9 assembly followed the protocol in Kroll et al., 2021. Cas9 protein (Alt-R S.p. Cas9 Nuclease V3, IDT, Cat. #1081058) was used to form RNP complexes. A gRNA targeting tyrosinase was included as a visual marker of CRISPR efficiency. The pooled Cas9/gRNA RNPs (tyr and target gene gRNAs) were injected into single-cell embryos.

#### List of designed crRNAs with their targeting exons

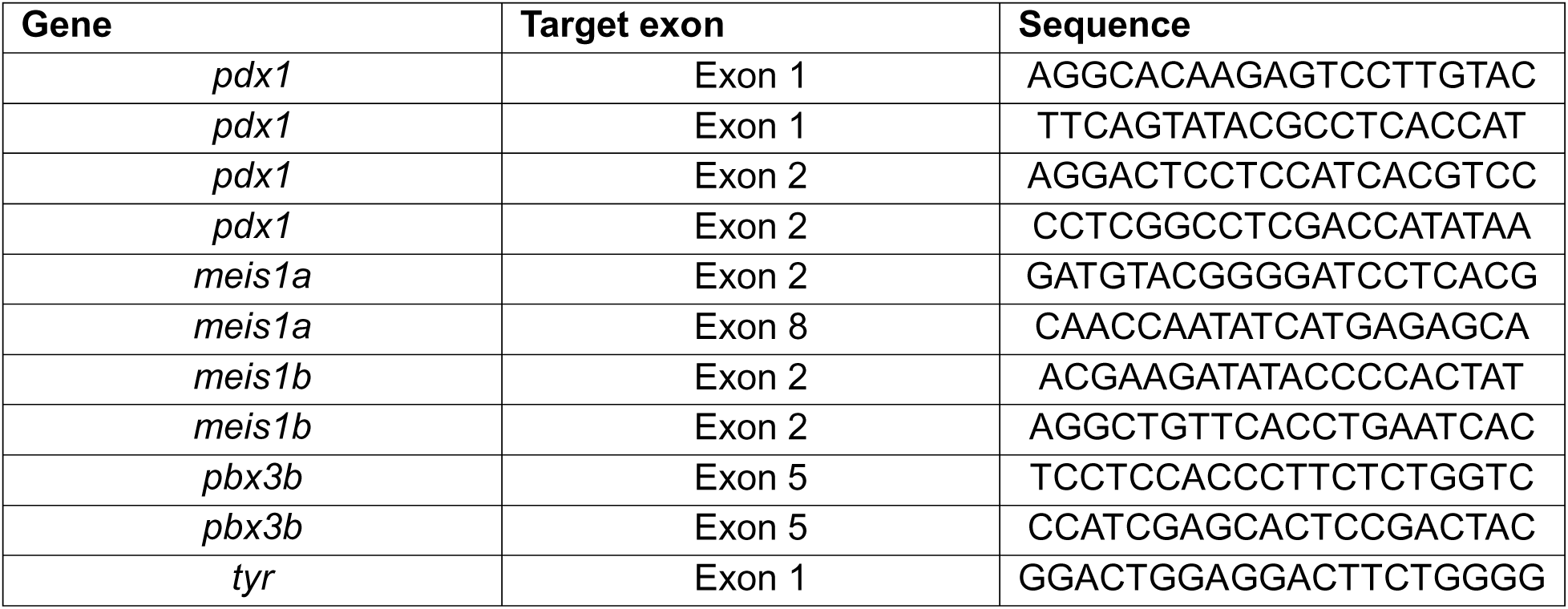

### Immunofluorescence and image acquisition

Whole-mount immunofluorescence was performed on 5dpf larvae fixed in 4% paraformaldehyde for 48 hours followed by skin removal from the abdomen region. The samples were permeabilized in 0.4% PBT (Triton-X-100) and blocked in 4% PBTB (BSA). Primary antibody, secondary antibody and DAPI staining were performed overnight at 4°C. Primary antibodies used in this study were anti-insulin (rabbit, GeneTex 128490) at 1:200 and anti-pdx1 (guinea pig, gift from Dr. Christopher Wright) at 1:200. Secondary antibodies at 1:500 dilutions used in this study were Alexa Fluor 568 anti-rabbit and Alexa Fluor 647 anti-guinea pig. Nile Red staining was performed at a concentration of 0.5ug/ml to quantify neutral lipid deposits. Samples were mounted in Vectashield and imaged using a Zeiss LSM 780 or Andor Dragonfly spinning disk confocal system (Andor Technology, Belfast, Northern Ireland). FIJI was used to add scale bars and PowerPoint was used for labeling.

### Quantification of bihormonal cells and lipid filled vacuoles in the islets

All quantifications were performed on z-stack confocal images of whole-mount transgenic zebrafish larvae at 5 dpf. *Ins*^+^/*sst1.1*^+^ cells were manually quantified as hybrid bihormonal cells and nile red+ spherical structures were manually quantified as neutral lipid filled vacuoles using Fiji (ImageJ 2.15.1/1.54f). Plotting and statistical analysis were performed using ordinary one-way ANOVA with Tukey’s multiple comparisons test in GraphPad (v11).

### MTZ ablation and pharmacological treatments at larval stages

Beta-cell ablation was induced by incubating 2.5 dpf larvae in 10 mM metronidazole (MTZ, Sigma-Aldrich, M3761) in E3 medium for 24 h. Larvae were thoroughly rinsed to remove MTZ and then incubated with the pharmacological compounds (GW6471, MedChemExpress, Catalog #: HY-15372) from the end of MTZ treatment until 5 dpf, with fresh compound solution replaced every 24 h.

### MTZ ablation in medaka

Ablation of beta cells in adult medaka was performed as described (Dasyani *et al*, 2019) with minor modifications: A 20 mM metronidazole solution was freshly prepared and adult fish were kept in 10 mM MTZ solution in fish system water for 24 h in the dark. Control fish were treated with fish system water containing 0.5% DMSO. After treatment, the MTZ was washed out with fish system water and the fish were transferred back to the husbandry system until the end of the experiment.

### Glucose quantification in medaka

For glucose quantification, brain tissues were harvested, snap-frozen in liquid nitrogen and then stored at −80L°C until further processing. Glucose measurements were determined using the BioVision Glucose Assay Kit (Cat #: ab65333) according to the manufacturer’s protocol. After thawing on ice, assay buffer was added at a ratio of 50 µL per 1 mg of brain tissue and the tissue samples were sonicated with an ultrasonic homogenizer (Bioruptor sonication device, Diagenode). Homogenates were then centrifuged at 13,000 × *g* and 50 µL of the supernatant was used for the glucose assay.

### Preparation of single-cell RNA-seq libraries

Suspensions of single cells of the zebrafish and medaka pancreas were performed as described in (Janjuha *et al*, 2018). Briefly, primary islets together with surrounding tissue were dissected and dissociated into single cells using TrypLE (Thermo Fisher Scientific, 12563029) supplemented with 0.1% Pluronic F-68 (Thermo Fisher Scientific, 24040032). Dissociation was carried out at 37 °C for 30 minutes at 450 rpm on a benchtop shaker. During dissection, β-cells were identified using fluorescence from Tg(ins:BB1.0L; cryaa:RFP) or Tg(ins:YFP-2A-NTR3, cryaa:mCherry) reporter lines to guide localization of the principal pancreatic islets. Following enzymatic digestion, TrypLE activity was inactivated by addition of 10% FBS. Cells were collected by centrifugation at 500 g for 10 min at 4°C, after which the supernatant was removed and the pellet resuspended in 500 µl HBSS lacking calcium and magnesium, supplemented with 0.1% Pluronic F-68. The suspension was filtered through a 30 µm cell strainer (Miltenyi Biotec, 130-041-407). Calcein violet (Thermo Fisher Scientific, C34858) was added to a final concentration of 1 µM, followed by incubation for 20 min at room temperature. Calcein-positive cells were isolated by fluorescence-activated cell sorting (FACS). Subsequent barcoding and library preparation were performed following the 10x Genomics single-cell RNA-seq protocol. Of note, this protocol led to lower detection of medaka endocrine pancreas cells compared to zebrafish, which limits the statistical power of the analysis shown in Figure 3G,H.

### Preparation of single-cell ATAC-seq libraries

Single-cell suspensions from zebrafish and medaka pancreas were prepared as described above. A total of 100,000 live cells were FACS-sorted into BSA-coated 1.5 mL tubes and pelleted by centrifugation at 500 g for 5 min at 4°C. Cells were washed once with chilled 1x PBS under the same centrifugation conditions.

For nuclei isolation, cell pellets were resuspended in 90 µL ice-cold lysis buffer (10 mM Tris-HCl pH 7.4, 10 mM NaCl, 3 mM MgCl₂, 0.1% Tween-20, 0.1% IGEPAL CA-630, 0.01% digitonin, 1% BSA) and gently pipetted. Lysis was performed on ice for 3 min. Subsequently, 100 µL wash buffer (10 mM Tris-HCl pH 7.4, 10 mM NaCl, 3 mM MgCl₂, 0.1% Tween-20, 1% BSA) was added, and nuclei were pelleted at 500 g for 5 min at 4 °C. The supernatant was carefully removed without disturbing the nuclei pellet.

The pellet was resuspended in 90 µL chilled 1x nuclei buffer (10x Genomics) and centrifuged at 500 g for 5 min at 4°C. After complete removal of the supernatant, nuclei were resuspended in 7 µL diluted nuclei buffer. Nuclei concentration was determined using trypan blue exclusion. Subsequent tagmentation, barcoding, and library preparation were performed following the 10x Genomics single-cell ATAC-seq protocol.

The sample preparation protocol for single-cell multiome libraries was largely the same as for scATAC-seq. In addition, we added 1mM DTT and 1U/ul of RNase inhibitor (stock 40U/ul) to lysis buffer, wash buffer and diluted nuclei buffer during preparation.

### Single-cell raw data processing and quality control

#### Read alignment and initial processing

Raw data of zebrafish pancreas scRNA-seq datasets GSE166052 (Singh *et al*, 2022) and GSE226841 (Mi *et al*, 2023), as well as new data generated here, were aligned to GRCz11 and processed using Cell Ranger v7.0.1 (10x Genomics) with default parameters. Ambient RNA contamination and putative doublets were removed using CellBender v0.3.2. Zebrafish scATAC-seq data were aligned to GRCz11 and processed using Cell Ranger ATAC v2.1.0 (10x Genomics) using default parameters. Zebrafish multiome datasets were aligned to GRCz11 and processed using Cell Ranger ARC v2.0.2 (10x Genomics) using default parameters. Medaka (Oryzias latipes) scATAC-seq data were aligned to the ASM223467v1 reference assembly and processed using Cell Ranger ATAC v2.1.0. Human multiome datasets were obtained from the PANCDB consortium (Kaestner *et al*, 2019) and aligned to the human reference genome hg38 using Cell Ranger ARC v2.0.2 (10x Genomics).

#### Processing and clustering of scRNA-seq data

Single cell RNA sequencing datasets were preprocessed using the Scanpy framework v1.9.5 (Wolf *et al*, 2018), and potential doublets and low-quality cells were filtered out. Single cell RNA datasets were merged and integrated using SCVI tools (Gayoso *et al*, 2022). Semi-supervised clustering was performed using the leiden algorithm based on the scVI latent space, which resulted in 25 clusters. These clusters were annotated using established marker genes. Subsets of the main endocrine islet populations were reclustered. Differentially expressed genes (DEGs) were identified for each cell type at different time points post ablation relative to the corresponding controls using Wilcoxon rank-sum test. Transcript counts per million (cpm) were calculated to obtain pseudobulk data per cell type and condition.

### Processing and integration of Bulk RNA data

FACS sorted data of islet cells pre and post ablation (GSE167187) was mapped to GRCz11 using STAR (Dobin *et al*, 2013) aligner and assigned to genomic features using featureCounts (Liao *et al*, 2014) We integrated our pseudobulk RNA data for cell type and conditions with more than 50,000 counts and annotated the FACS sorted data based on transcriptional similarity using PCA clustering (Figure S1C).

### Processing and clustering of scATAC-seq and paired sequencing data

Single cell ATAC and paired multiome datasets were processed using the Signac (Stuart *et al*, 2021) framework. Potential doublets and low-quality cells were filtered out. Peaks were called on all fragments of high-quality cells using MACS2 (Zhang *et al*, 2008). Gene activity scores were calculated based on Signac’s GeneActivity function, which includes counts in the proximity of 2 kb of TSS and protein coding regions of annotated genes. The peak count matrix was normalized using term frequency inverse document frequency (TF-IDF) and dimensions were reduced using singular value decomposition (SVD). Harmony (Korsunsky *et al*, 2019) was used for dataset integration, followed by clustering with the Louvain algorithm. 27 clusters were identified and annotated by established marker gene activity scores.

Endocrine cell populations were identified using the canonical marker genes, and coverage tracks of fragments overlapping marker-gene loci were plotted using Signac’s CoveragePlot function. Displayed fragments covered annotated genes as well as 2 kb upstream and downstream regions. To include weaker signals, peak calling was repeated. We ran MACS2 on sets of fragments for each cell type and timepoint. To create a uniform peak set, we then merged overlapping or directly adjacent peaks into a non-redundant peakset. For obtaining pseudo-bulk data, counts per million (cpm) were calculated per cell type and condition. We ran normalization, dimensional reduction, integration and clustering as described before. For the multiome dataset, paired gene expression data of *sst1.1* and *ins* were mapped on the dimensional reduction.

### Differential accessibility analysis and dataset balancing

To reduce biases from unequal cell numbers, we applied a two-step subsampling strategy. Cells were first downsampled to achieve comparable representation across datasets, followed by subsampling within combined cell type and treatment groups to obtain similar group sizes. Rare populations of gip and ε-cells could not be fully balanced due to low abundance.

Differentially accessible peaks were identified using logistic regression as implemented in Signac (‘FindMarkers’), including read depth as a latent variable. Pairwise comparisons were performed across all groups, and peaks were considered significant at adjusted p ≤ 0.01 and log_2_ fold-change ≥ 1. Next, we filtered for the most accessible peaks (top 20%) of grouped cell types and calculated pairwise Pearson’s correlations across groups.

### Multiome analysis and differential accessibility

Multiome δ1-cells were classified into *ins_high_*, *ins_mid_*, and *ins_low_* categories based on normalized *ins* expression (defined as >1 SD above, within, or <1 SD below the mean, respectively), and differential peak accessibility was tested using logistic regression. Multiome coverage tracks were plotted as described above.

### Motif Accessibility

The JASPAR 2024 Vertebrate database was used to extract position weight matrices (PWMs) for TF binding (Rauluseviciute *et al*, 2024). Zebrafish orthologue gene names were annotated using ENSEMBL (Dyer *et al*, 2025) and OMA (Altenhoff *et al*, 2024) databases. The peak set was then scanned for accessible motifs using motifmatchr (Schep, 2025) and chromVAR (Schep *et al*, 2017) was used to score variability in motif accessibility between the cells. This analysis was performed for both the full scATAC-sequencing dataset and the endocrine subset. PCA and UMAP were then run on chromVAR scores for data visualization.

### Gene regulatory network analysis

To identify gene regulatory networks, we adapted a workflow previously published by (Zhu *et al*, 2023), as described below:

#### Input pseudobulk data and indexing

Cell types and treatment conditions were pseudo-bulked as described above. Data from 2-7 dpa were grouped as the early response, and data from 14–21 dpa as the late response. Pseudobulk RNA data were filtered to include only genes with a minimum of 5 cpm and pseudobulk ATAC data were filtered to include only peaks with a minimum count of 3 cpm. ATAC peaks were annotated relative to genes using zebrafish genome annotations. Transcription start sites (TSS) were defined based on the first exon of each transcript. Peaks were then classified as candidate cis-regulatory elements (cCREs) based on genomic proximity: promoter cCREs (±2 kb from TSS), proximal cCREs (±25 kb), and distal cCREs. Promoter and proximal cCREs were associated with the nearest gene. To identify distal cCREs, we used Cicero to detect regions with correlated accessibility across cells while applying a distance-dependent penalty, with a maximum distance of ±500 kb. Distal cCREs were defined as cCREs that exhibited significant co-accessibility with promoter peaks (co-accessibility score ≥ 0.1). (Pliner *et al*, 2018) cCRE-gene links were merged into a table and duplicates were removed prioritizing promoter > proximal > distal assignments.

#### Calculation of cCRE-gene links

cCRE–gene links assigned in the previous step were evaluated by calculating Pearson correlations between cCRE accessibility and expression of the corresponding putative target gene across pseudobulk samples. To generate a background distribution, RNA and ATAC sample columns were randomly shuffled and gene–cCRE assignments were independently permuted. Correlations were recomputed on the perturbed data, and observed and permuted correlations were combined for downstream comparison. A correlation cutoff was then determined for each position to achieve a false discovery rate (FDR) of below 0.15 (Figure S10A-C).(Pliner *et al*, 2018)

#### Calculation of TF-cCRE links

Putative transcription factor (TF)–cCRE regulatory links were inferred by integrating motif annotations with filtered cCRE–gene connections. Only cCREs passing the correlation thresholds required for a FDR of 0.15 were retained, using position-specific cutoffs for promoter (R>0.5), proximal (R>0.61), and distal elements (R>0.46). TF–cCRE pairs were defined based on motif occurrences within cCREs, and correlations between TF expression and cCRE accessibility were computed across pseudobulk samples using Pearson correlation. To estimate a null distribution, TF–cCRE assignments were randomly permuted and sample orders of RNA and ATAC vectors were independently shuffled prior to correlation calculation. Observed and perturbed correlations were compared to assess the strength of TF–cCRE regulatory associations. A correlation cutoff (R>0.7) was then determined to achieve a false discovery rate (FDR) of below 0.15 (Figure S10D).

#### TF modules in differentially expressed genes

Differentially expressed genes for each cell type and condition were considered significant if they showed log fold change > 1 and adjusted p < 0.01. Significant DEGs were mapped to their associated cCREs using the previously defined cCRE–gene connections, generating sets of regulatory elements linked to each DEG group. Enrichment was calculated using Fisher’s exact test comparing the frequency of TF motif–containing cCREs within each DEG-associated cCRE set against the background of all analyzed cCREs. Resulting p-values were corrected for multiple testing using the Benjamini–Hochberg method.

For visualization, the top enriched TFs per cluster were selected and plotted as a dot plot, where dot size represents the enrichment odds ratio and color indicates the significance of enrichment (−log10 adjusted p-value).

### GO term enrichment analysis

For each TF, orthologues of the corresponding target-gene sets were analyzed for pathway enrichment using Enrichr v3.4 against the *WikiPathways 2023 Human* (Agrawal *et al*, 2024) gene set library. For each TF, the top five pathways were selected based on adjusted p-value ranking.

### Usage of LLMs

ChatGPT (OpenAI, GPT-5.3) was used to improve clarity of text and assist in code refinement. All outputs were critically reviewed and validated by the authors.

