## supplementary figures for "Pre-existing chromatin accessibility primes δ-cells for injury-induced endocrine plasticity"

### Supplementary Figure 1

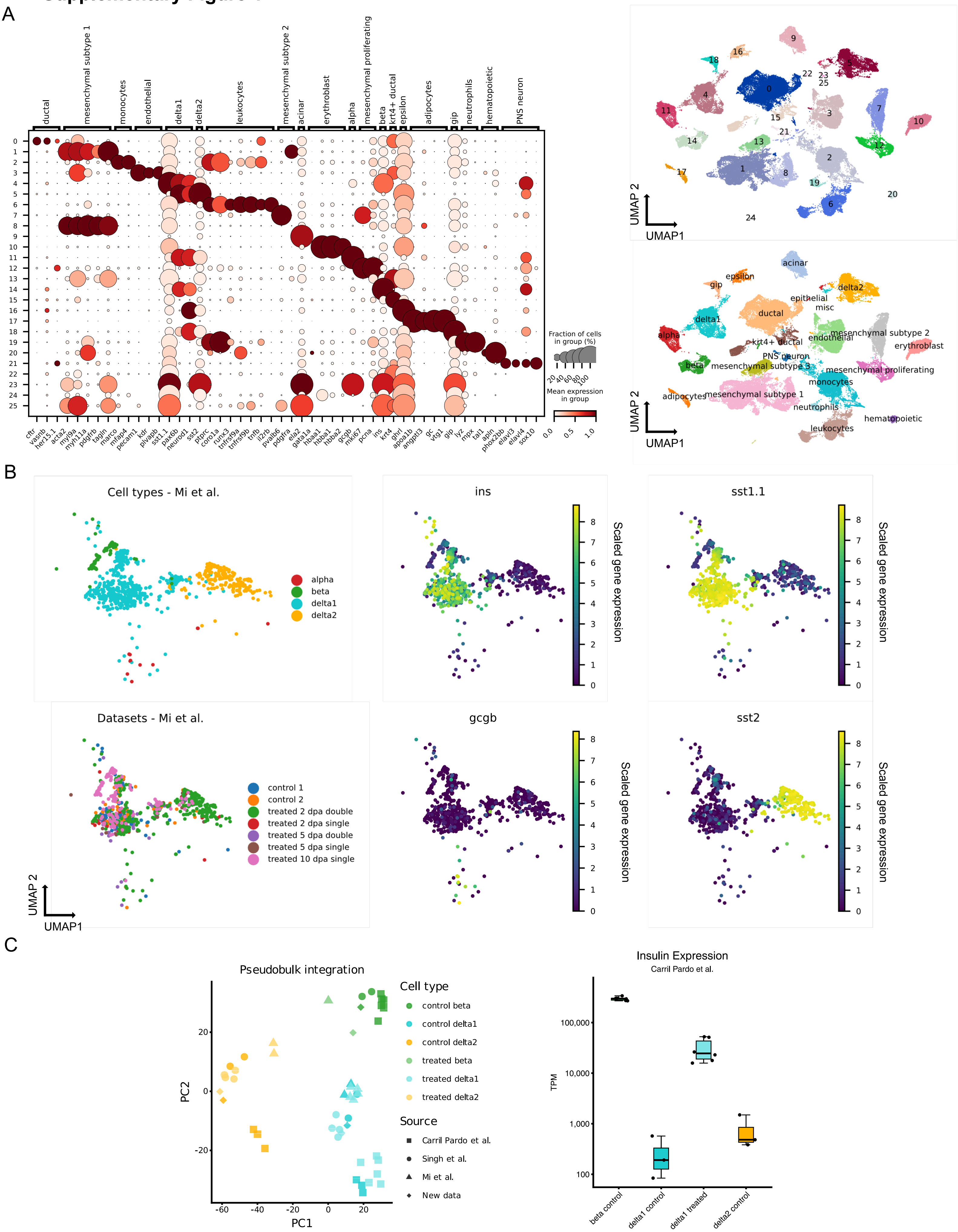

**Supplementary Figure 1: A single-cell atlas of β-cell ablation shows that primed δ1-cells provide early insulin restoration across datasets in adult zebrafish**

A) Cluster annotation of the integrated scRNA-sequencing dataset and utilized marker genes for annotation. All differential genes per cluster are listed in Supplementary Dataset S2**.**

B) Sub clustering of **Figure 1D** showing only data from Mi et al. 2023.

C) PCA integration of FACS sorted bulk RNA-seq data (Carril Pardo *et al*, 2022) and pseudo bulked scRNA-seq data. Gene expression of *ins* in bulk RNA-sequencing data categorized by pseudobulk similarity.

### Supplementary Figure 2

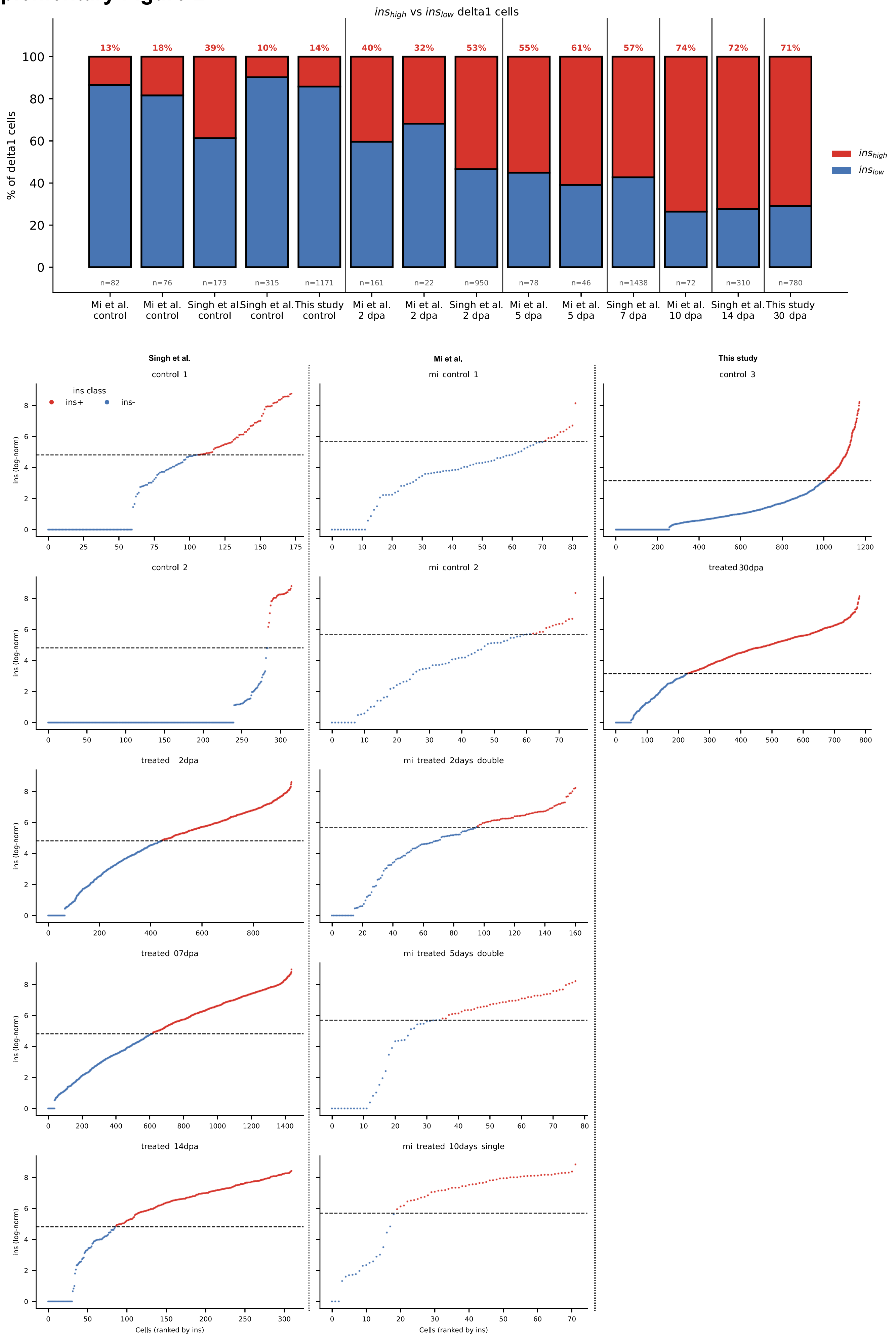

**Supplementary Figure 2: Up to 74% of δ1-cells turn *ins*$^+$ after β-cell loss in adult zebrafish.**

Classification of δ1-cells from different sources into *ins*$^+$ and *ins*$^-$. Cutoffs were set as +1SD of the mean *ins* expression of the respective controls. Bars show the percentage of *ins*$^+$ δ1-cells over time post ablation.

#### Supplementary Figure 3

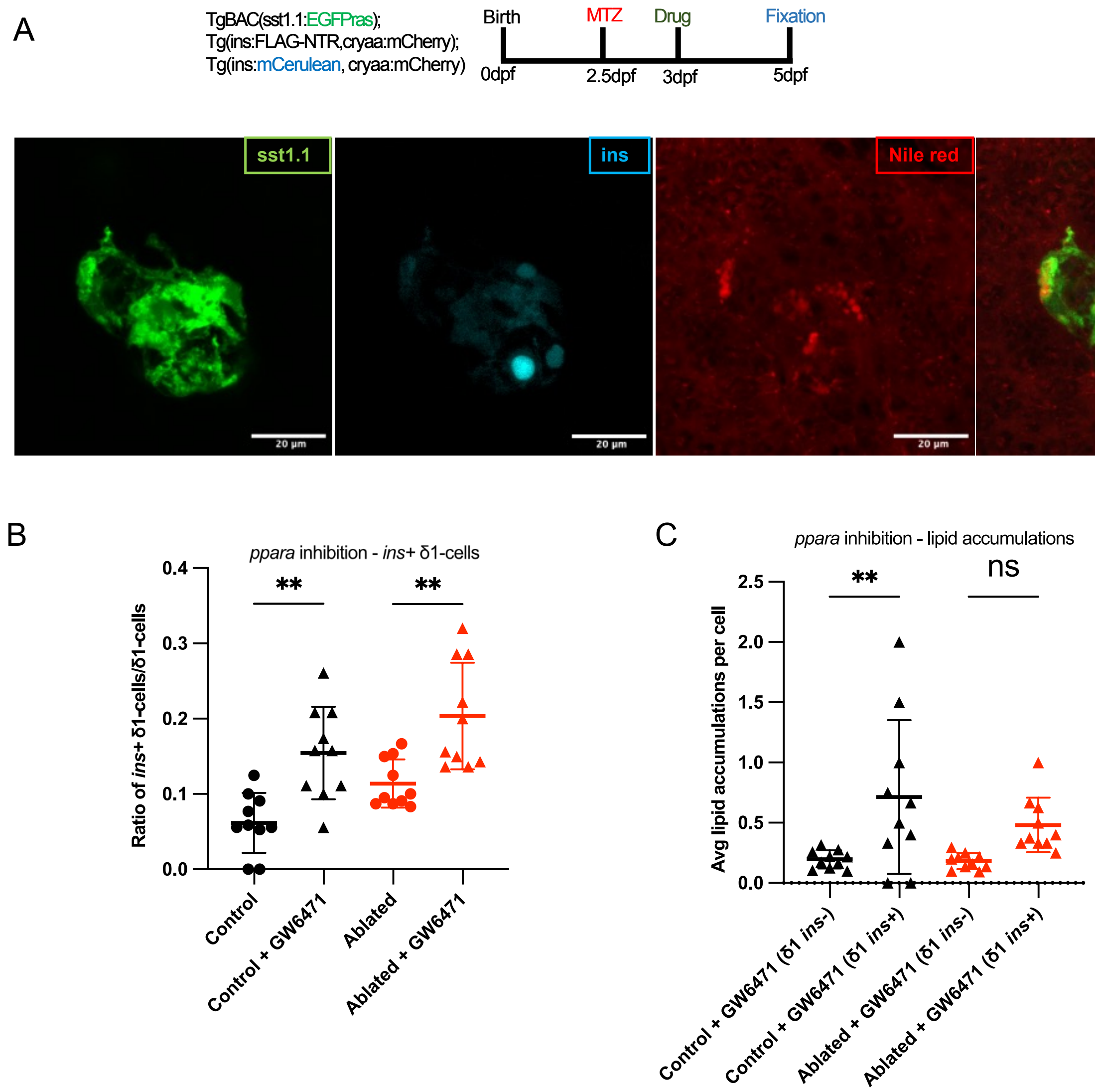

**Supplementary Figure 3: *PPARα* inhibition promotes *ins*$^+$ δ1-cell formation in zebrafish larvae**

A) Representative image of a larval pancreatic islet post β-cell ablation with lipid accumulations in δ1-cells. δ1-cells are labelled by sst1.1:EGFP-Ras (green), insulin-expressing cells by ins:NLS-mCerulean (cyan), and neutral lipids by Nile Red (red). Scale bar: 20 μm.

B) Pharmacological inhibition of *PPARα* mediated fatty acid oxidation significantly increases the number of *ins*$^+$ δ1-cells, following β-cell ablation as well as in the unablated DMSO control. Larvae were treated with the *PPARα* antagonist GW6471. Each point represents one islet (n = 10 larvae). Statistical significance was determined using one-way ANOVA with Tukey's multiple comparisons test. Bars indicate mean ±1 SD.

C) Quantification of lipid accumulations in δ1-cells after treatment with the *PPARα* antagonist GW6471. Inhibition reduces the prevalence of lipid accumulations in *ins*$^+$ δ1-cells, in both treated and untreated larvae. Each point represents one islet (n = 10 larvae). Statistical significance was determined using one-way ANOVA with Tukey's multiple comparisons test. Bars indicate mean ±1 SD.

### Supplementary Figure 4

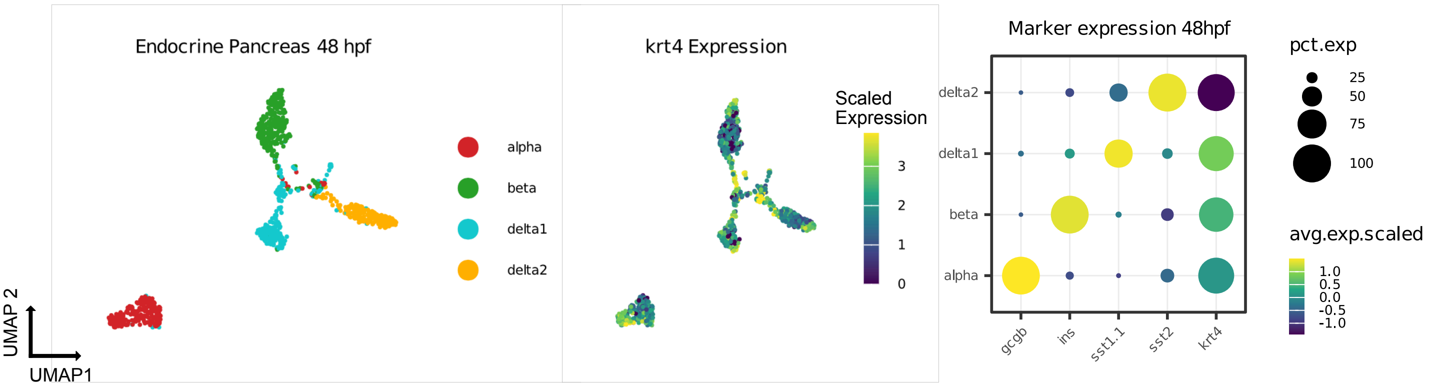

**Supplementary Figure 4: δ1-cells are *krt4* positive during zebrafish development**

scRNA-seq data from zebrafish islets at 48 hpf show that endocrine cells are partially positive for expression of the ductal marker *krt4*.

### Supplementary Figure 5

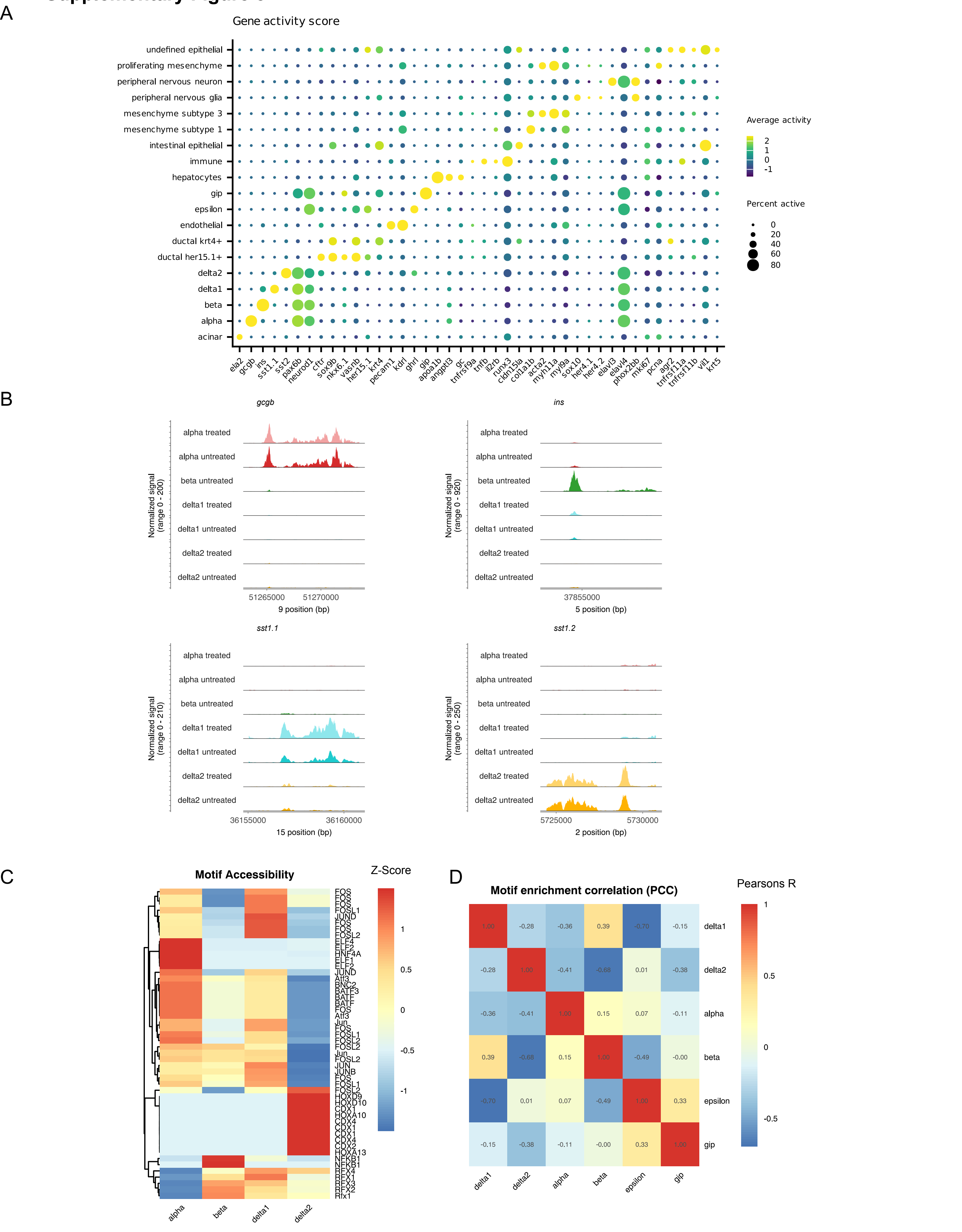

**Supplementary Figure 5: Single cell ATAC sequencing data show chromatin accessibility landscape of the zebrafish pancreas.**

A) Marker gene activity used for annotation of zebrafish scATAC-seq data.

B) Chromatin accessibility plots of endocrine marker genes pre and post β-cell ablation.

C) Top motif accessibility by zebrafish cell type.

D) Pearson correlation coefficient of motif accessibility by cell type.

### Supplementary Figure 6

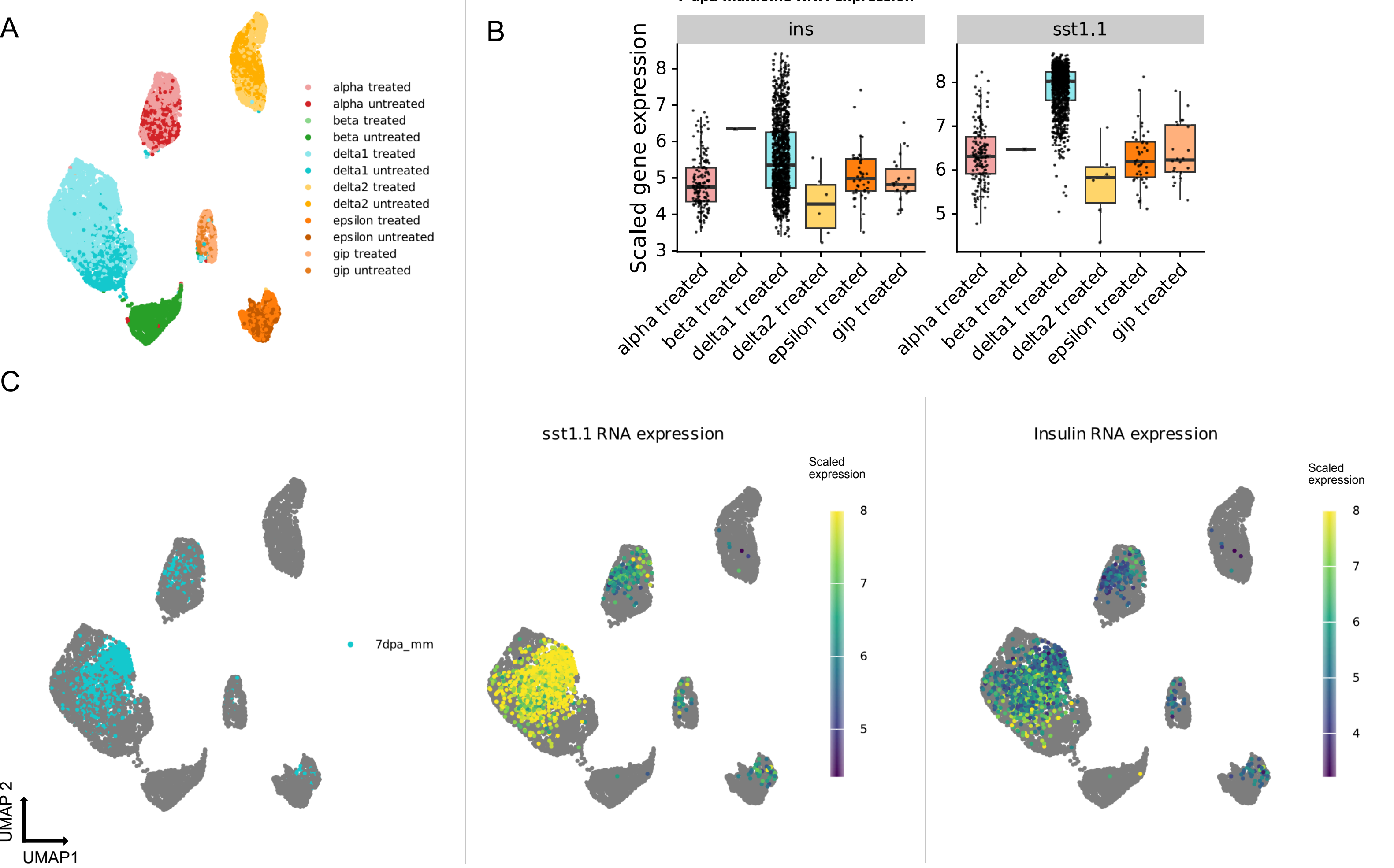

**Supplementary Figure 6: Paired scRNA/scATAC-seq multiome data show δ1-cells express insulin without major chromatin remodeling in zebrafish.**

A) UMAP representation of scATAC-seq data of all endocrine cells (similar to Figure 2D).

B) Paired gene expression of *sst1.1* and *ins* in 7 dpa multiome data classified by scATAC-seq modality.

C) scATAC-seq data clustering does not separate *ins*$^{+}$ and *ins*$^{-}$ δ1-cells.

### Supplementary Figure 7

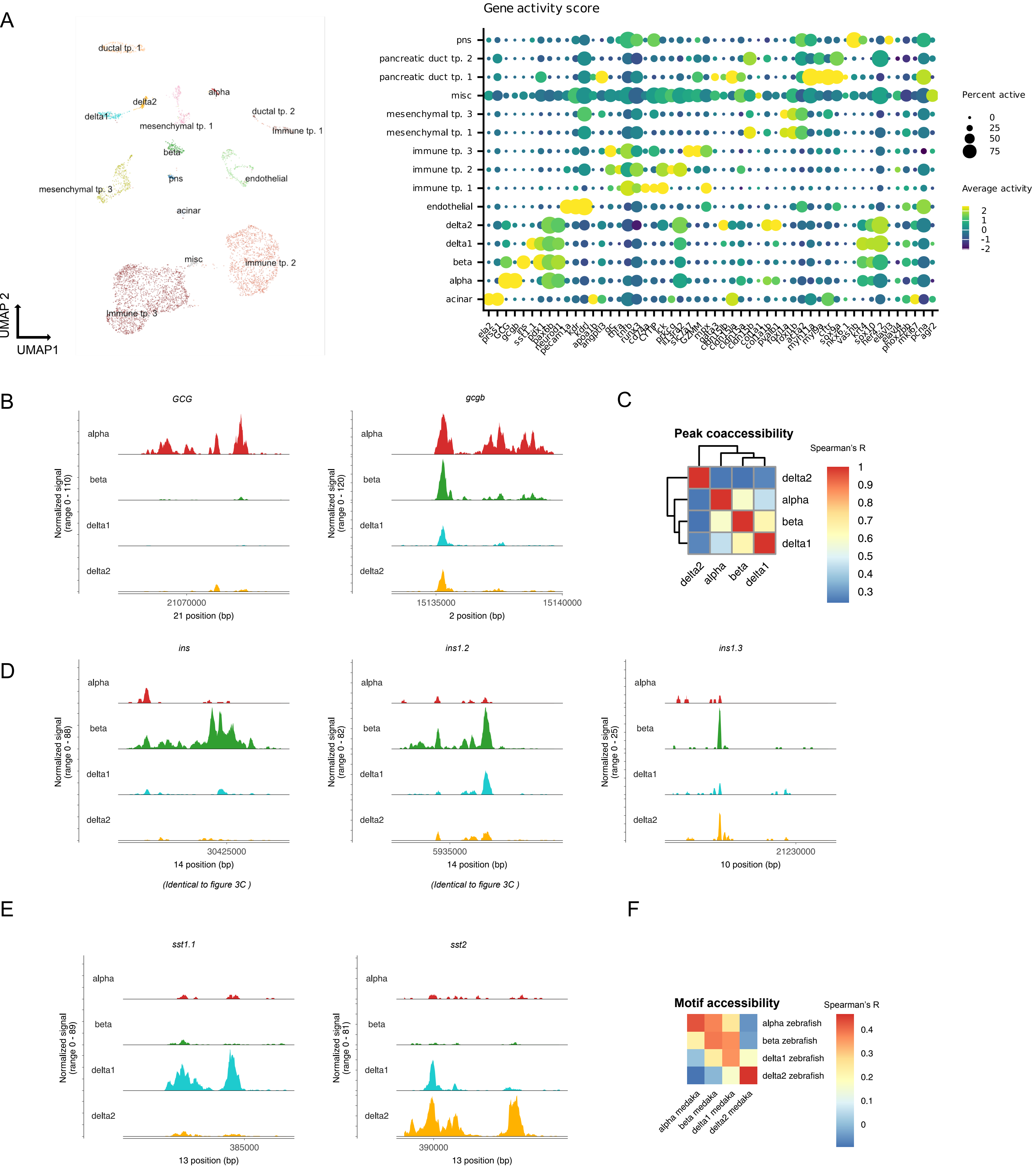

**Supplementary Figure 7: Chromatin accessibility landscape of the medaka pancreas.**

A) Whole scATAC-seq dataset of medaka pancreas and surrounding tissues and gene accessibility used for cell type annotation. All differential accessible genes per cluster are listed in Supplementary Dataset S7**.**

B) Gene accessibility of α-cell markers *GCG* and *gcgb*.

C) Pearson correlation of peak co-accessibility in medaka islets.

D) Gene accessibility of medaka *ins* isoforms.

E) Gene accessibility of medaka *sst* isoforms.

F) Interspecies co-accessibility of motifs by cell type (Spearman’s correlation coefficient)

#### Supplementary Figure 8

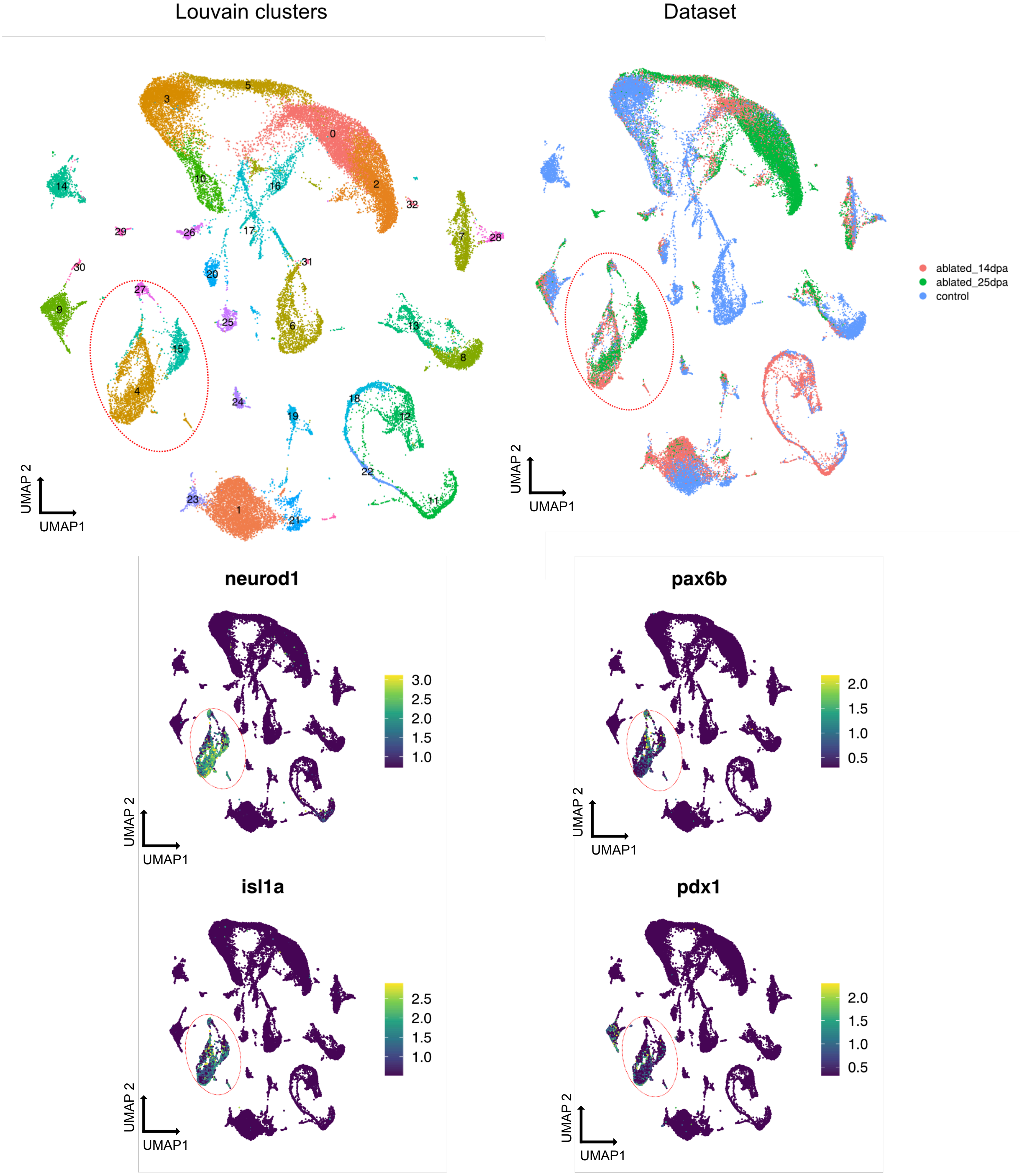

**Supplementary Figure 8: Time course scRNA sequencing data pre and post β-cell ablation of the medaka pancreas.** Samples were taken from control animals and at 14 and 25 days post ablation. Louvain cluster 4, 15 and 27 were identified as endocrine populations based on TF expression (*pax6b, pdx1, neurod1, isl1a*).

### Supplementary Figure 9

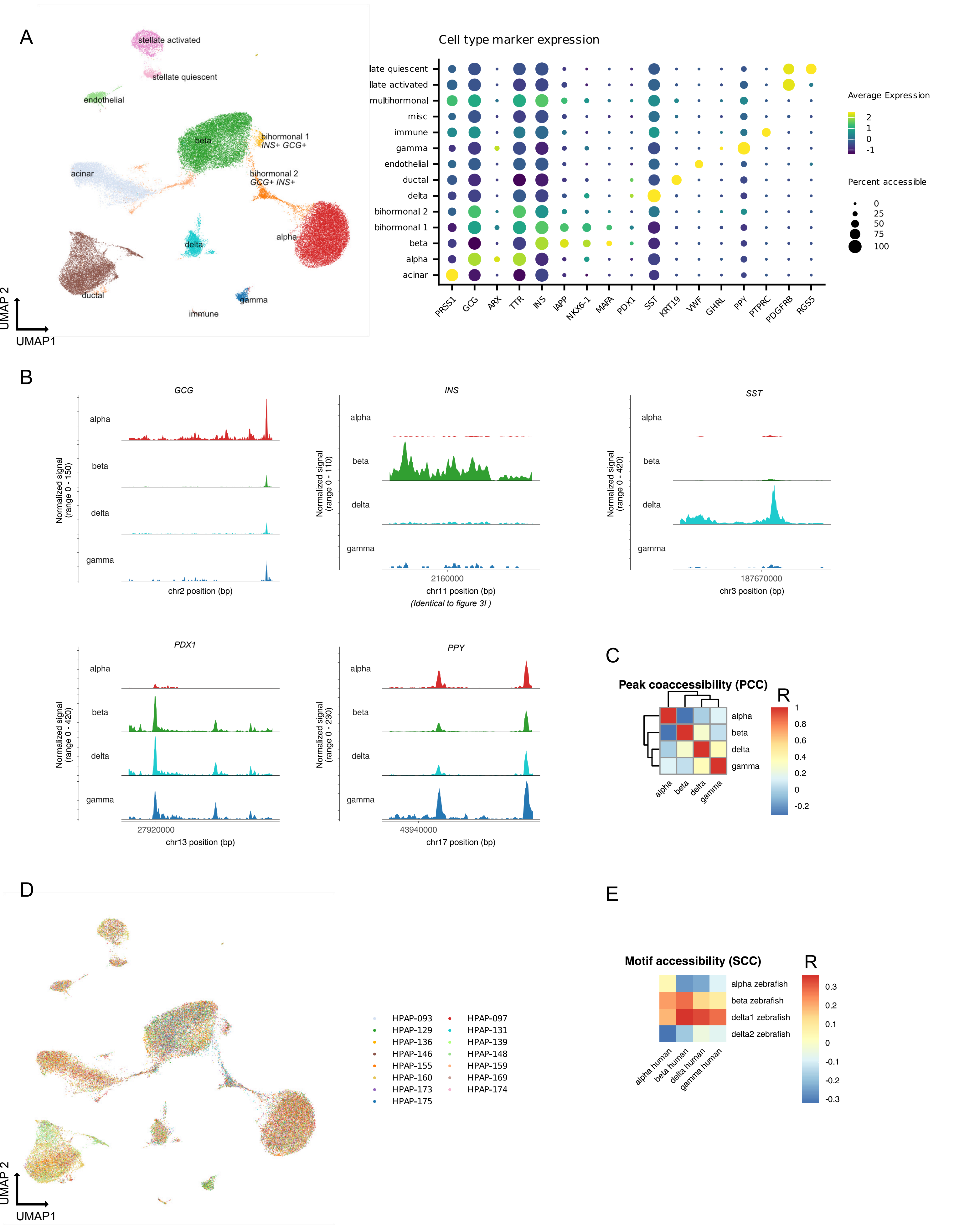

**Supplementary Figure 9: Chromatin accessibility landscape of the healthy human pancreas**

A) UMAP representation and paired marker gene expression of multiome data from pancreata of 16 non-diabetic individuals as listed in Supplementary Dataset S1.

B) Chromatin accessibility by cell type of endocrine marker genes.

C) Pearson's correlation coefficient of peak co-accessibility.

D) Samples included in the dataset.

E) Interspecies coaccessibility of motifs by cell type (Spearman’s correlation coefficient)

### Supplementary Figure 10

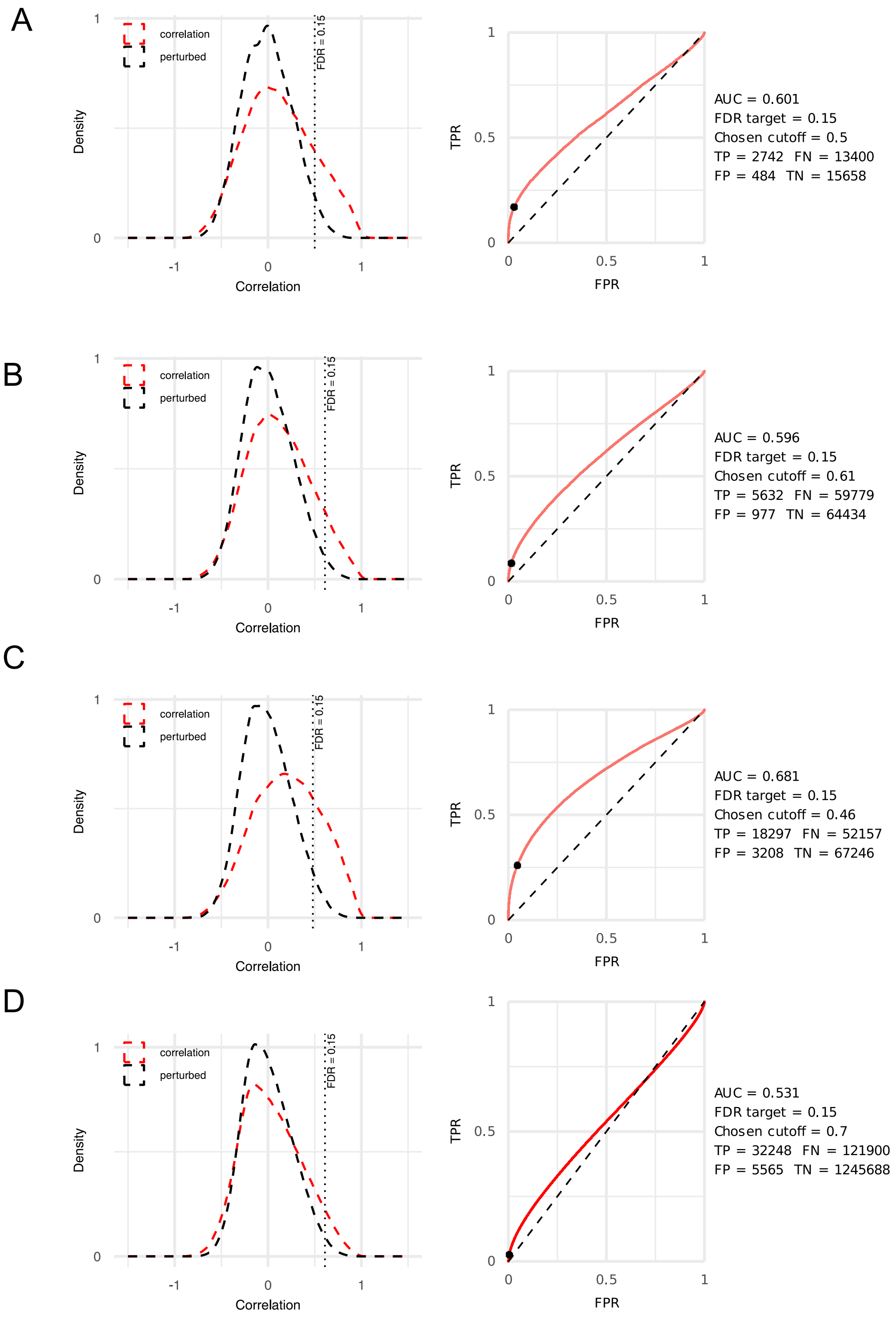

**Supplementary Figure 10: Quality control and cutoff settings for GRN processing.**

A) Pearson correlation of promoter (± 2kb) cCRE accessibility to target gene expression (red) vs random (black).

B) Pearson correlation of proximal (± 25kb) cCRE accessibility to target gene expression (red) vs random (black).

C) Pearson correlation of Cicero selected distal (± 500kb) cCRE accessibility to target gene expression (red) vs random (black).

D) Pearson correlation of transcription factor gene expression to putative target cCREs (red) vs random (black).

### Supplementary Figure 11

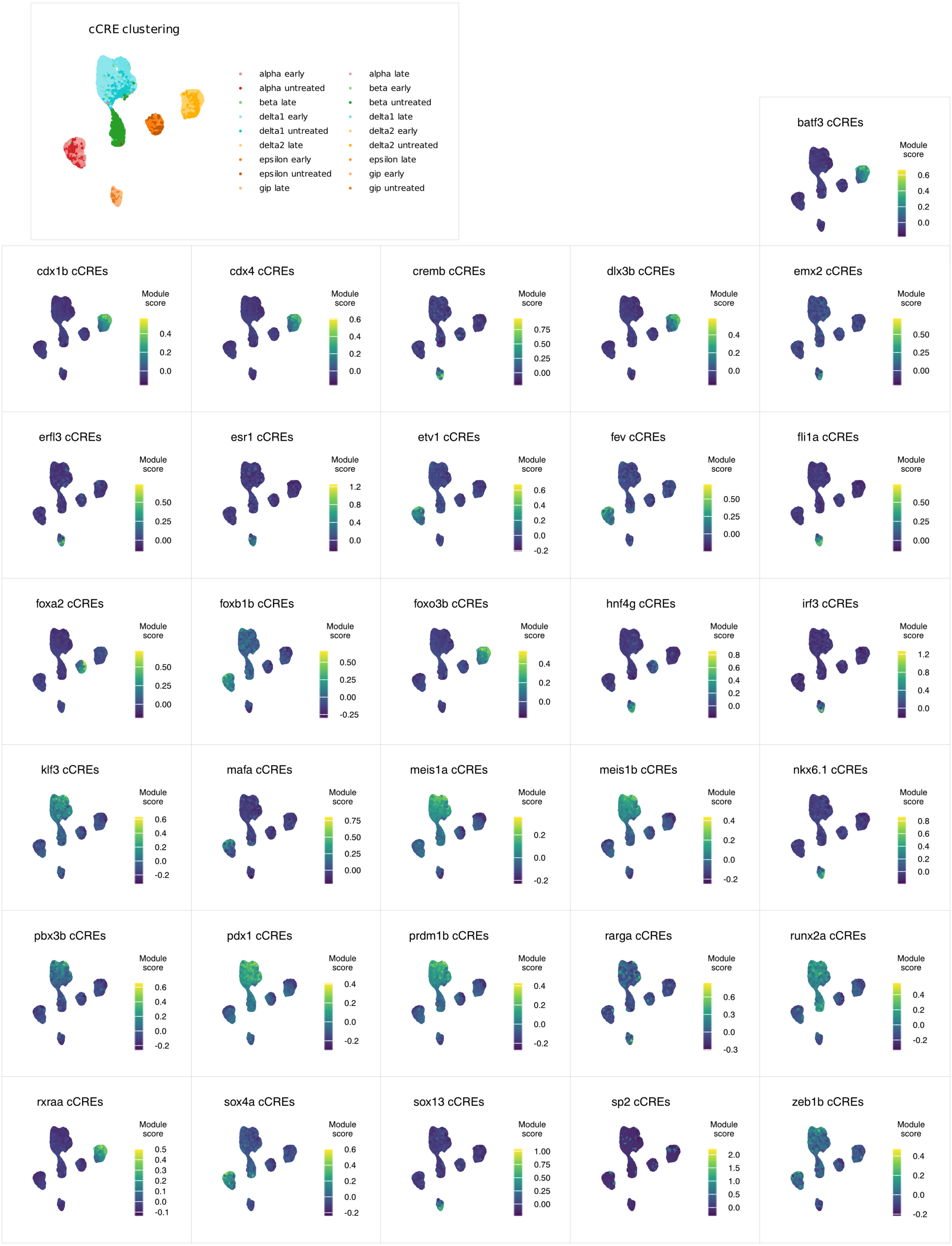

**Supplementary Figure 11: TF module accessibility in endocrine cells**

UMAP representation of endocrine cells using only cCREs which are linked to TF modules. Module accessibility scores represent the mean accessibility of all cCREs which are linked to a specific transcription factor, which allows identification of cell type specific programs.

### Supplementary Figure 12

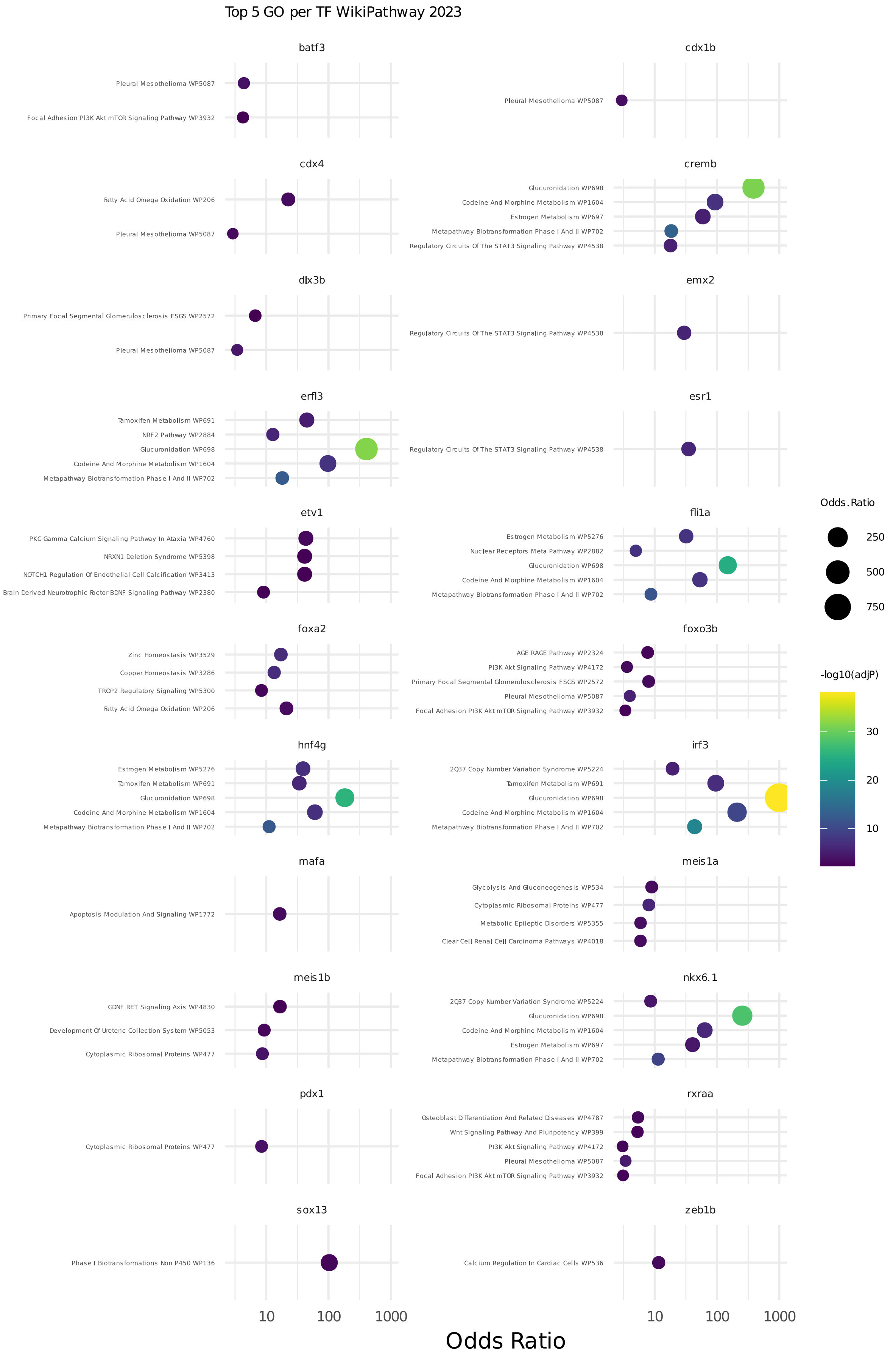

**Supplementary Figure 12: Gene ontology enrichment terms of transcription factor target genes**

Gene ontologies of differentially expressed genes post ablation linked to the respective transcription factors. Enrichment analysis was performed using enrichR with the WikiPathway database 2023.

### Supplementary Figure 13

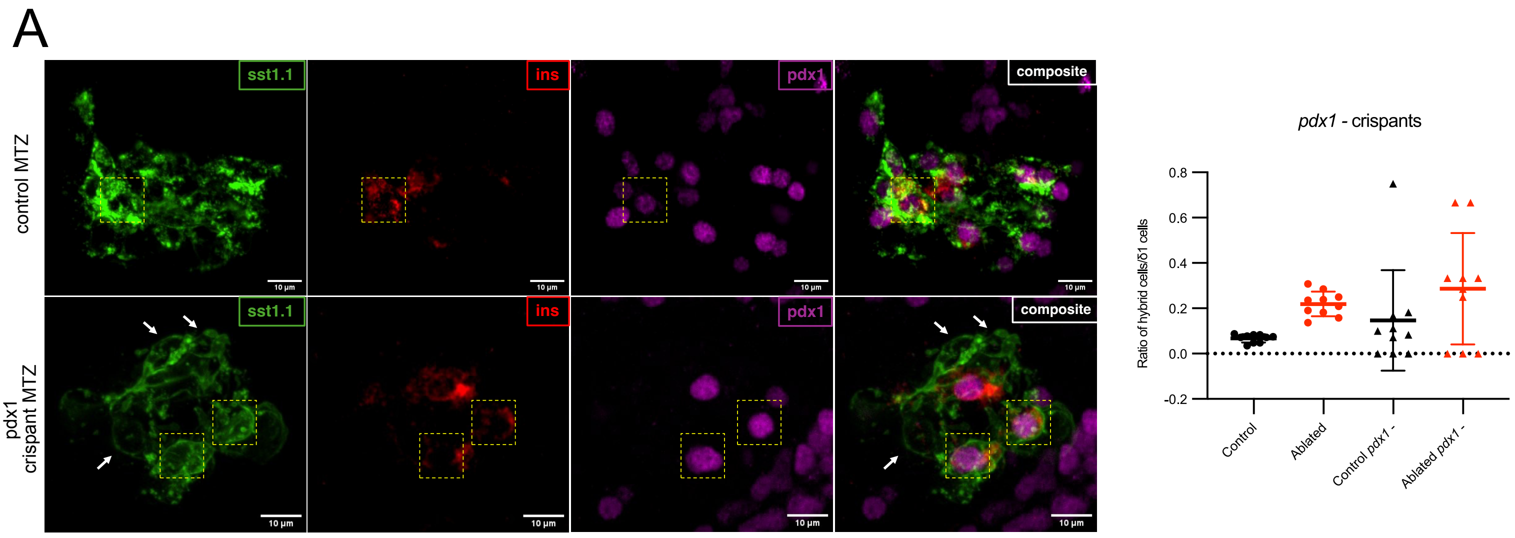


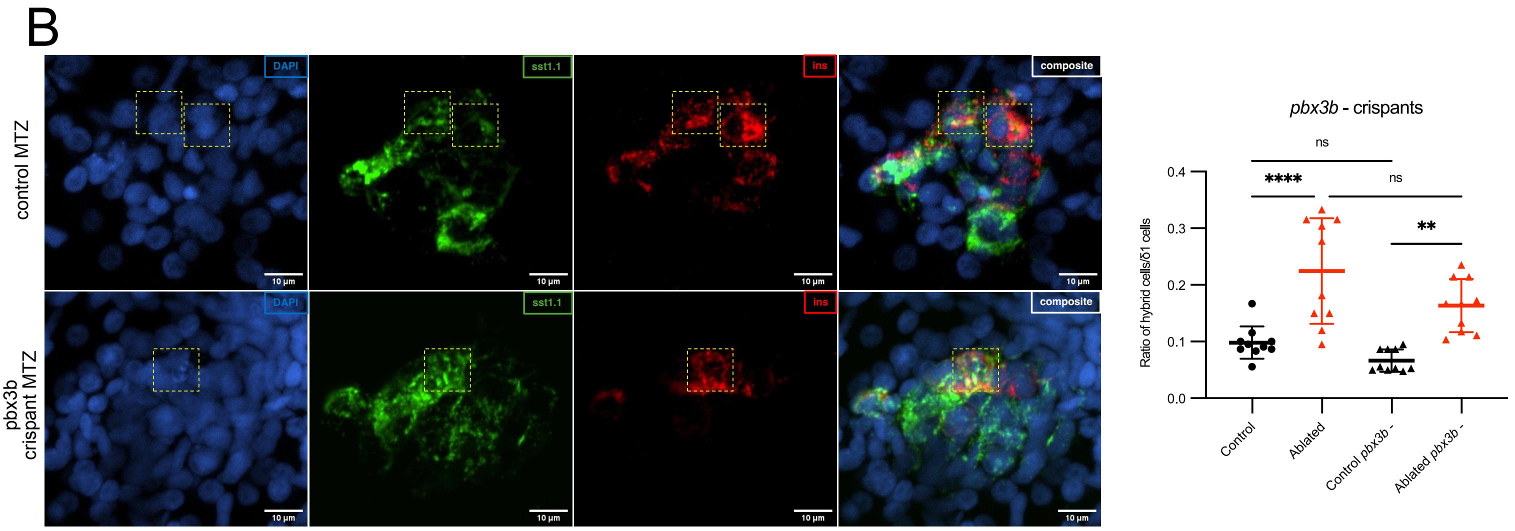

**Supplementary Figure 13: Crispants of *pdx1* show disrupted islets while perturbations of *pbx3b* do not reduce numbers of hybrid cells in zebrafish larvae.**

A) Representative confocal images and quantifications of hybrid cells in pancreatic islets after β-cell ablation and *pdx1* F0 knockouts. In control larvae, δ1-cells (*sst1.1*$^{+}$) activate insulin expression and form bihormonal hybrid cells. In F0 knockouts of *pdx1* targeting exon 2, islets are mostly disrupted. Antibody staining against insulin (red) and *pdx1* (magenta) show that existing hybrid cells are *pdx1* positive. White arrows indicate δ1-cells negative for pdx1 expression and yellow dotted squares indicate *sst1.1*$^{+}$/*ins*$^{+}$ double-positive cells.

B) Representative confocal images and quantifications of hybrid cells in pancreatic islets after β-cell ablation and *pbx3b* F0 knockouts with antibody staining against insulin (red). In control larvae, δ1-cells (*sst1.1*$^{+}$) activate insulin expression and form bihormonal hybrid cells. In F0 knockouts of *pbx3b*, the reduction of hybrid cells is not statistically significant. Yellow dotted squares indicate *sst1.1*$^{+}$/*ins*$^{+}$ double-positive cells.
